# Natural variation of immune epitopes reveals intrabacterial antagonism

**DOI:** 10.1101/2023.09.21.558511

**Authors:** Danielle M. Stevens, Alba Moreno-Pérez, Alexandra J. Weisberg, Charis Ramsing, Judith Fliegmann, Ning Zhang, Melanie Madrigal, Gregory Martin, Adam Steinbrenner, Georg Felix, Gitta Coaker

## Abstract

Plants and animals detect biomolecules termed Microbe-Associated Molecular Patterns (MAMPs) and induce immunity. Agricultural production is severely impacted by pathogens which can be controlled by transferring immune receptors. However, most studies use a single MAMP epitope and the impact of diverse multi-copy MAMPs on immune induction is unknown. Here we characterized the epitope landscape from five proteinaceous MAMPs across 4,228 plant-associated bacterial genomes. Despite the diversity sampled, natural variation was constrained and experimentally testable. Immune perception in both *Arabidopsis* and tomato depended on both epitope sequence and copy number variation. For example, Elongation Factor Tu is predominantly single copy and 92% of its epitopes are immunogenic. Conversely, 99.9% of bacterial genomes contain multiple Cold Shock Proteins and 46% carry a non-immunogenic form. We uncovered a new mechanism for immune evasion, intrabacterial antagonism, where a non-immunogenic Cold Shock Protein blocks perception of immunogenic forms encoded in the same genome. These data will lay the foundation for immune receptor deployment and engineering based on natural variation.

**Significance Statement:** Plants recognize pathogens as non-self using innate immune receptors. Receptors on the cell surface can recognize amino acid epitopes present in pathogen proteins. Despite many papers investigating receptor signaling, the vast majority use a single epitope. Here, we analyzed the natural variation across five different epitopes and experimentally characterized their perception in plants. We highlight the importance of analyzing all epitope copies within a pathogen genome. Through genetic and biochemical analyses, we revealed a mechanism for immune evasion, intrabacterial antagonism, where a non-immunogenic epitope blocks perception of immunogenic forms encoded in a single genome. These data can directly inform disease control strategies by enabling prediction of receptor utility and deployment for current and emerging pathogens.

## Introduction

Plants contain hundreds of innate immune receptors capable of recognizing diverse pathogens. Biotic organisms carry microbe-associated molecular patterns (MAMPs), fragments of larger biomolecules such as proteins, carbohydrates, and lipids, that are recognized by surface-localized pattern recognition receptors (PRRs) (1-3). PRRs include receptor-like kinases (RLKs) and receptor-like proteins (RLPs) that recognize conserved MAMPs or damage-associated molecular patterns, resulting in PRR-triggered immunity (PTI) (3,4). Activation of PTI induces multiple defense responses including: the production of reactive oxygen species (ROS) and ethylene, apoplast alkalization, activation of mitogen-activated protein kinase (MAPK) cascades, transcriptional reprogramming, and callose deposition at the cell wall culminating in disease resistance (4,5).

The most well-studied PRR is FLAGELLIN-SENSING 2 (FLS2), which recognizes the 22-amino acid epitope, flg22, from bacterial flagellin FliC (6-9). Unlike most PRRs, *FLS2* is conserved throughout the plant kingdom (7,9,10). Since the flg22-FLS2 discovery, researchers have uncovered other epitope-receptor pairs restricted to certain plant families. For instance, the Elongation Factor Tu (EF-Tu)-derived epitope elf18 interacts with the EFR receptor, which is present in the Brassicaceae family (11,12). The 22-amino acid epitope of cold shock protein (CSP), csp22, interacts with CORE and a second flagellin epitope flgII-28 interacts with FLAGELLIN-SENSING 3 (FLS3), both solanaceous RLKs (13-16). Finally, the necrosis-and-ethylene inducing peptide 1 (Nep1)-like protein (NLP) epitope nlp20 interacts with RLP23 found in Brassicaceae (17-19). These receptors recognize certain bacterial MAMPs, including those present in pathogens and commensals.

Proteins that carry MAMP sequences are thought to be important for microbial survival and fitness. For example, bacterial flagellin enables bacterial swimming and swarming and is critical for the colonization of certain hosts (20). Different regions of flagellin monomer, FliC, are recognized by different receptors in plants and mammals despite the flagellin apparatus requiring multiple components (21). CSPs are an ancient protein family first described in *Escherichia coli* with roles in RNA chaperoning in cold environments (22). This can include unwinding RNA, maintaining RNA stability, limiting internal cleavage, and enhancing expression via antitermination. Recent genetic work on other bacterial systems has shown additional post-translational regulatory roles related to virulence, stress tolerance, pili formation, and biofilm development (23-28). CSPs are composed of two critical motifs, RNP-1 and RNP-2, near the N-terminus of the small beta-barrel like structure that binds broadly to nucleic acids (22). While the regulatory function of CSPs appear conserved across bacteria, their targets and sequences are highly variable. NLPs are cross-kingdom phytotoxic virulence factors (17,19). Flagellin, CSPs, and NLPs are considered expendable, though their loss is predicted to confer fitness costs. Conversely, EF-Tu is a highly abundant, essential protein that transports aminoacyl-tRNAs to the ribosome during translation (29). MAMP-encoded gene function and abundance likely have an impact on their evolution in the context of plant immune interactions.

Using flg22 perception as a model, three general epitope outcomes have been described: immune activation, evasion, and antagonism. Many Gram-negative plant pathogens carry immunogenic epitopes, though some can evade perception (7,30-32). MAMP evasion, also known as masking, occurs by accumulating sequence variation that prevents epitope binding (33,34). MAMPs can act antagonistically and block subsequent perception of immunogenic epitopes from other bacteria by binding to the primary receptor and inhibiting proper signaling complex formation (34,35). Sequence variation within the perceived epitope is thought to be constrained by protein functionality, though this has only been studied in the flg22 epitope and FliC protein (8). How polymorphism affects protein function versus immune perception has not been investigated for other MAMP-encoded proteins. In addition, how MAMP copy number variation (CNV) encoded within a single genome impacts immune outcomes remains elusive. Therefore, many questions remain regarding how natural epitope variation interplays with protein function and plant immune perception.

To broadly understand how natural evolution impacts immune outcomes across five different MAMPs, we mined 34,262 MAMP epitopes from 4,228 whole bacterial genomes. Each MAMP displayed substantial copy number and sequence variation. While the theoretical number of MAMP variants is astronomically large, natural variation is constricted, making it experimentally testable to characterize MAMP evolution. We focused on characterizing the immunogenic outcomes of the EF-Tu (elf18) and CSP (csp22) MAMPs for sequence and CNV. Elf18 displays minimal sequence variation and gene expansion, with most variants inducing strong immune responses. In contrast, csp22 displays considerable variation in epitope sequence, CNV, and immune outcomes. Using a combination of phylogenetics, genetics, and biochemistry, we revealed conserved non-immunogenic CSPs in a subset of bacterial genera, some of which antagonize perception of immunogenic forms encoded in the same genome. We then characterized an actinobacterial CSP that acts as an intrabacterial antagonist of the CORE receptor, which enables immune evasion.

## Results

### Evolutionary trajectories depend on the immunogenic feature and bacterial genera

Here, we sought to assess the presence and diversity of bacterial proteinaceous MAMPs. Across 13 genera, we identified plant-associated bacterial genomes with representing members from alpha-, beta-, and gamma-proteobacteria as well as Gram-positive actinobacteria (Dataset S1). We extracted features from the following proteins and their corresponding MAMP epitopes: bacterial flagellin (flg22 and flgII-28), cold-shock protein (csp22), EF-Tu (elf18), and Nep1-like protein (nlp20). A computational pipeline was developed that extracts peptides using a modified BlastP protocol and local protein alignment, polymorphic ends and off-target correction, and clonality filtering (Fig. 1A). This enabled mining of epitopes in a gene description-dependent and - independent manner allowing for comprehensive genome-derived epitope comparisons (SI Appendix, Table S1).

**Fig. 1.**
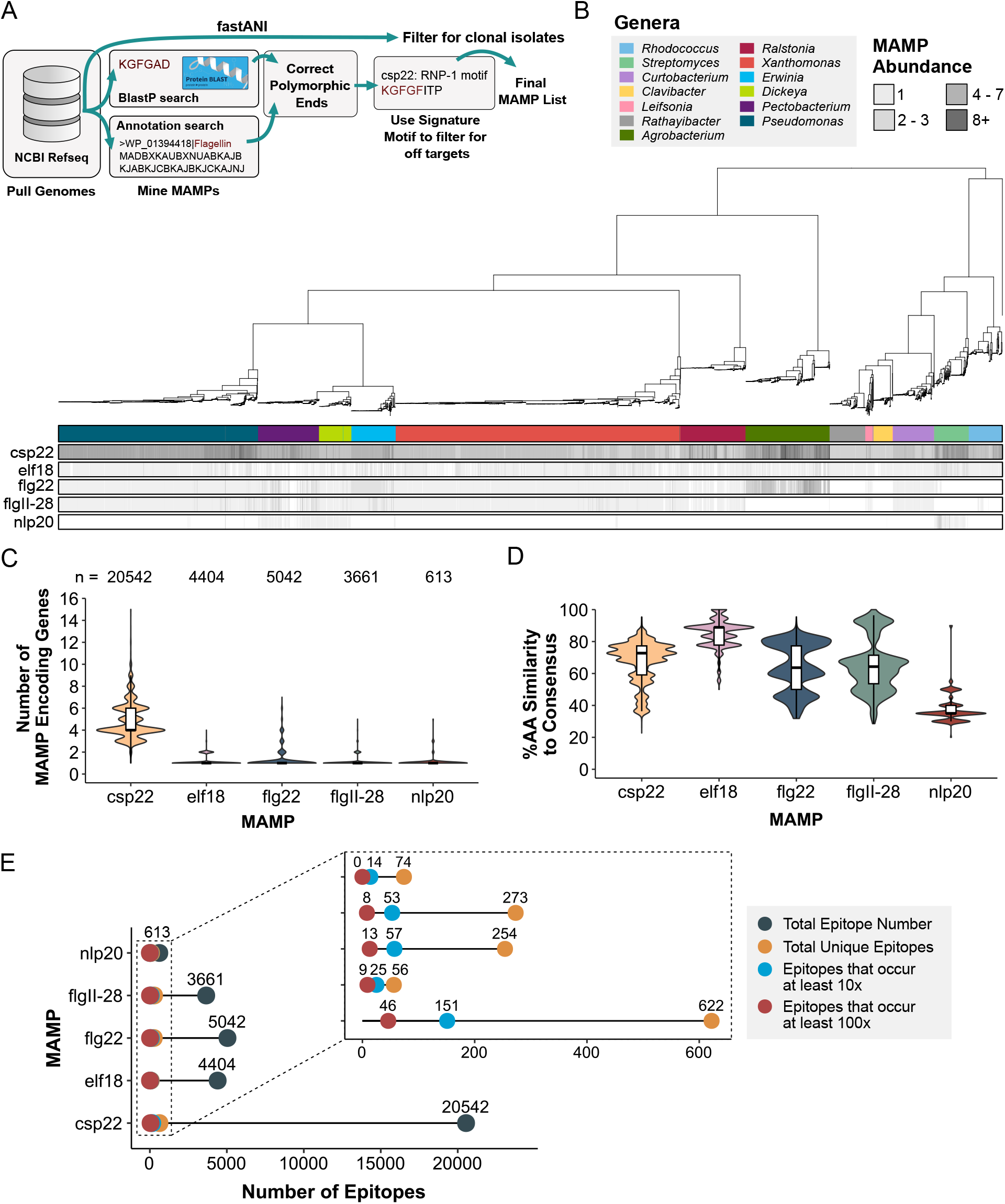
Epitopes from diverse plant-associated bacteria exhibit different evolutionary trajectories. (*A*) Pipeline to mine for MAMP epitopes from 4,228 plant-associated bacterial genomes (see Methods Section for details). (*B*) Maximum-likelihood phylogeny built from 74 housekeeping genes with tips labeled by genera. Each bar below presents the number of epitopes (csp22, elf18, flg22, flgII-28, nlp20) present in each genome. (*C*) A violin plot showing the number of MAMP-encoded genes from each genome separated by MAMP type. Tukey’s boxplots are plotted on top. The number of epitopes assessed are listed at the top. (*D*) A violin plot of percent amino acid (AA) similarity of each epitope variant in comparison to each respective consensus sequence across all bacteria sampled. Tukey’s boxplots are plotted on top. (*E*) Lollipop plot displaying the number of unique MAMP epitopes and their occurring frequencies.

Differences in sequence and CNV were detected in a MAMP-dependent manner. From the 4,228 genomes, 34,262 epitopes were extracted and their abundance was plotted on a 74 gene maximum-likelihood tree (Fig. 1B). Notably, there are different patterns of epitope CNV in a lineage-specific manner (Fig. 1B, SI Appendix, Fig. S1A). The MAMPs elf18, flg22, flgII-28, and nlp20 are primarily encoded by single copy genes, whereas csp22 variants displayed expansion and are encoded by 1 to 15 genes (Fig. 1C, SI Appendix, Fig. S1A). Next, we investigated CNV across 13 genera (SI Appendix, Fig. S1A). Strains had an average of four CSPs, with variation in a genera and species-dependent manner. For example, CSP CNV was increased in *Agrobacterium* and *Streptomyces* (average of eight and seven paralogs, respectively) compared to *Dickeya, Pectobacterium, Xanthomonas, Ralstonia, Curtobacterium, Rathayibacter*, and *Clavibacter* (average of three to five copies) (SI Appendix, Fig. S1A). Although *fliC* is primarily a single copy gene in 80% (2921/3631) of the genomes, additional copies can be observed in *Erwinia, Pectobacterium*, and *Agrobacterium* (SI Appendix, Fig. S1A). Similarly, EF-Tu is primarily a single copy gene in 84% (2825/3346) of analyzed genomes, with additional copies predominantly found in Gram-negative bacteria apart from *Streptomyces* (Fig. 1B-C, SI Appendix, Fig. S1A). Nlp20 was only found in 13% (565/4228) of the genomes and predominantly in necrotic bacteria such as *Streptomyces, Pectobacterium*, and *Dickeya* (SI Appendix, Fig. S3A-B).

We investigated MAMP conservation compared to the ‘consensus’ sequence commonly used in the literature for immune assays. When all genera were analyzed together, each MAMP exhibited different distributions compared to the consensus epitope (Fig. 1D). The cumulative distributions of flg22 and flgII-28 variants closely mirror each other and are derived from different regions of the same gene, FliC (Fig. 1D, SI Appendix, Fig. S1B, Fig. S3C). However, a lower number of flgII-28 epitopes were detected (3661) compared to flg22 (5042), likely due to higher sequence diversity in the flgII-28 region (S1 Appendix, Fig S2). Elf18 and nlp20 variants are the most and least conserved, respectively (Fig. 1D, SI Appendix, Fig. S3C).

When analyzing MAMP distributions in individual genera, interesting trends are identified (SI Appendix, Fig. S1A). Diversification of each FliC epitope in Gram-negative bacteria does not always mirror each other. *Dickeya* and *Pectobacterium* exhibit similar amino acid similarity and CNV for both flg22 and flgII-28 (SI Appendix, Fig S1A). However, *Agrobacterium* has a multimodal distribution for flg22 and unimodal distribution for flgII-28 (SI Appendix, Fig S1A). Many Gram-positives exhibit similar csp22 diversification, indicating ancient CSP paralog emergence (SI Appendix, Fig. S1A). We also compared epitope variation independent of a ‘consensus’ sequence by calculating amino acid similarity in an all-by-all manner (SI Appendix, Fig. S1B). These results were complementary to the comparison to the ‘consensus’ epitope, with different patterns of diversity for each MAMP. Overall, we observed that epitope abundance and sequence diversification evolved in both a MAMP-based and genera-derived manner.

Genes that encode MAMP epitopes are postulated to be either essential or conditionally-essential for bacterial survival. Considering their ancient origin, we assessed the total degree of epitope variation, which we found was constrained. Except for nlp20, the total number of MAMP epitopes detected ranged from 3,661 to 20,542 (Fig. 1E). However, the total number of unique epitope variants was much lower with csp22 exhibiting the highest number (622) and elf18 displaying the lowest number (56, Fig. 1E). When considering the frequency of epitope variants, there was substantially fewer that occurred at least 10 or 100 times (between 8 and 151, Fig. 1E). Collectively, these data indicate it is possible to experimentally test the impact of natural variation on immune outcomes across thousands of bacteria.

### Most elf18 variants are immunogenic

To characterize the functional diversity of elf18, 25 variants were synthesized and assessed for their immunogenicity using ROS, an early, quantitative output of immune induction (5,35). ROS production by NADPH oxidases is required for biological processes, stress responses, and the induction of systematic acquired resistance, providing broad-spectrum resistance (36). The 25 elf18 epitopes (100 nM) were screened for their ability to induce ROS on *Arabidopsis thaliana* Col-0 and the *EFR* receptor mutant line. Water and the *E. coli-*derived ‘consensus’ elf18 epitope were used as controls (SI Appendix, Fig. S4A). Of the 25 variants tested, 76% (19/25) displayed equal ROS output in comparison to the consensus, 16% were weakly immunogenic (4/25), and 8% (2/25) were non-immunogenic (SI Appendix, Fig. S4A). No measurable ROS was produced in the *efr* mutant line (SI Appendix, Fig. S4A). To confirm our ROS screen was effective in classifying immunogenicity, a blinded, independent evaluation displayed the same conclusions as the initial screen (SI Appendix, Fig. S4B). Interestingly, all non-immunogenic variants and half of the weakly immunogenic variants were derived from a second EF-Tu locus in *Streptomyces*, that clustered separately (Fig. 2A). While all epitopes were derived from annotated EF-Tu loci, the second *Streptomyces* EF-Tu copy displayed the most divergent elf18 sequence (Fig. 2A, SI Appendix, Fig. S4A).

**Fig. 2.**
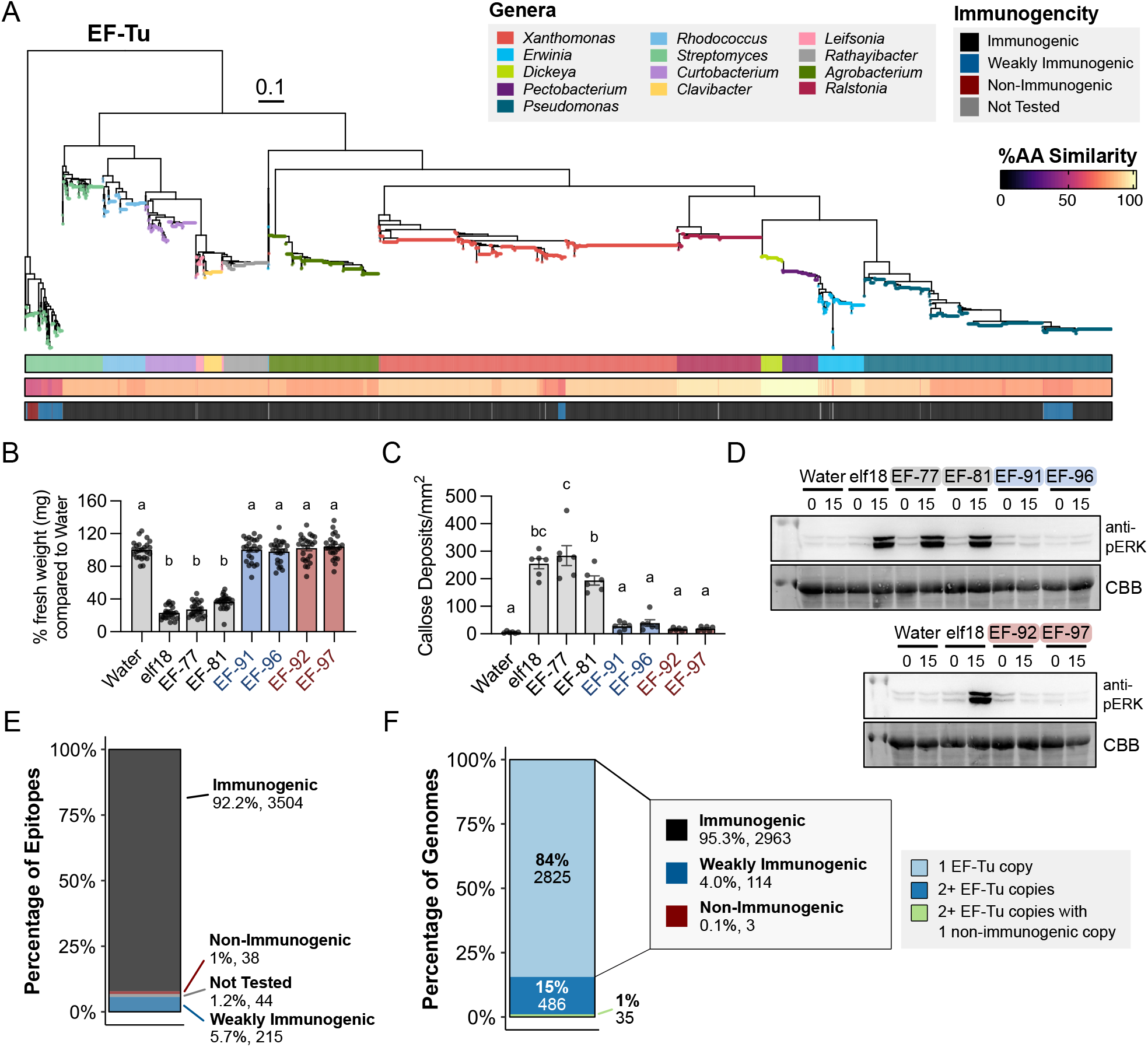
elf18 exhibits minimal diversification and most variants are immunogenic. (*A*) Maximum-likelihood phylogenetic tree of EF-Tu. Tips were colored by genera, second bar represents percent amino acid similarity to consensus, and third bar indicates immunogenicity by ROS from Supplemental Figure 4. (*B*) *Arabidopsis* seedling growth inhibition after treatment with water or elf18 peptides (100 nM). One point = 1 plant. Eight plants per biological replicate. A one-way ANOVA and Tukey’s mean comparison was used to determine significance, p<0.0001. (*C*) Quantified callose deposition in Col-0 after treatment with water and elf18 variants (1 μM). Values are from one representative experiment which includes an average of at least two images per leaf from two leaves per plant, three plants per treatment. Statistical analyses were performed as described in (*B*), p<0.0001. (*D*) Induction of MAPK by elf18 variants (100 nM) in Col-0 at zero- and 15-minutes post infiltration. CBB = protein loading. All experiments were repeated at least three times with similar results. (*E*) Summary of immunogenic outcomes of EF-Tu-encoded elf18 epitopes from Supplemental Figure 4. (*F*) Summary of immunogenic outcomes with respect to EF-Tu copy number.

Next, we characterized mid-to late-stage immune outputs including MAPK induction (100 nM), callose deposition (1 uM), and seedling growth inhibition (100 nM) for a subset of epitopes that exhibited different ROS immunogenicity. All secondary immune outputs were congruent with each other. All immune outputs by any elf18 epitope variant were abolished in the *efr* mutant (SI Appendix, Fig. S4C-E). Epitopes EF-77 and EF-81, immunogenic by ROS, inhibited seedling growth, induced MAPK activation, and displayed high amounts of callose deposition (Fig. 2B-D). In contrast, the weakly immunogenic and non-immunogenic variants EF-91, 92, 96, and 97 failed to induce mid-to late-stage immune responses, mirroring water or mock controls (Fig. 2B-D). Elf18 variants EF-91 and EF-96 can uncouple their immune outputs, potentially enabling pathogen evasion of strong immune response. Similarly, EF-80 is weakly immunogenic by ROS and has been previously shown to not induce robust callose deposition, a late-stage immune output (5,37). To understand the relationship between immunogenic outcomes and protein evolution, we developed a maximum-likelihood tree of EF-Tu and plotted genera, the percent amino acid similarity of each epitope to the consensus, and immunogenicity by ROS (Fig. 2A). The EF-Tu tree structure was indicative of taxonomic origin (Fig. 2A). The 25 experimentally tested epitope variants enabled the determination of immunogenicity of 98.8% (3757/3801) of elf18 variants across 3346 plant-associated bacteria (Fig. 2E). Most elf18 encoded variants (92.2%, 3504/3801) induce strong immunity when *EFR* was present (Fig. 2E). Furthermore, of the genomes that encode for more than one EF-Tu locus, only 35 have one immunogenic copy and second non-immunogenic copy (Fig 2F). While immune outcomes of weakly-immunogenic variants from *Streptomyces, Xanthomonas*, and *Pseudomonas* are comparable, their polymorphisms are unique and their respective EF-Tu proteins do not cluster phylogenetically, indicating convergent evolution (Fig. 2A, SI Appendix, Fig. S4A) (37). Across all elf18 variants, polymorphisms were position-dependent, showcasing epitope variation may be constrained by EF-Tu function (SI Appendix, Fig. S4A).

### Diversification of CSPs contributes to differential immunogenic responses

Unlike EF-Tu and elf18, CSP and csp22 exhibited CNV and epitope diversification (Fig. 1C-D, SI Appendix, Fig. S2A). This provided an opportunity to characterize the functional diversity of a second MAMP with a distinct evolutionary trajectory. Therefore, 65 csp22 epitope variants (200 nM) were screened for their ability to induce ROS on Rio Grande tomatoes. Water and the *Micrococcus luetus-*derived ‘consensus’ csp22 epitope were used as controls (SI Appendix, Fig. S6A) (13). We tested all variants with one of two independent *core-*deficient lines developed in the Rio Grande cultivar via CRISPR-cas9 (SI Appendix, Fig. S5 and S6A). Each *core* line failed to perceive csp22 but could still perceive flg22 and flgII-28 (SI Appendix, Fig. S5B). Across all 65 variants, no ROS was produced in the *core* mutant line, demonstrating *CORE* specificity (SI Appendix, Fig. S6A). Only 54% (35/65) of csp22 epitope variants displayed equal ROS output in comparison to the consensus, while 11% (7/65) were weakly immunogenic and 25% (23/65) were non-immunogenic (SI Appendix, Fig. S6A). A subset of the csp22 variants (25) were also assessed for their ability to induce ethylene production (1 uM). Though the resolution was qualitative, most epitope variants were consistent with our ROS screen (SI Appendix, Fig. S7).

Csp22 variation was explored across diverse CSP-domain loci. The CSP domain ranges from 65-75 amino acids in length and carries RPN-1 and RPN-2 nucleic acid binding motifs (22). While the CSP domain was conserved, considerable diversity in gene sequence length was found (SI Appendix, Fig. S6B). We identified CSPs carrying extra domains including calcium binding and DUF domains (SI Appendix, Fig. S6B). We identified an *Agrobacterium* CSP containing two cold shock protein domains fused together with a linker (Fig. S6B). Classical CSPs (<75 amino acids, no additional domains) predominantly encoded immunogenic epitopes. However, several weakly- or non-immunogenic epitopes were encoded within CSPs with unique domain architecture (SI Appendix, Fig. S6). Some CSPs were conserved across multiple genera, while others were restricted to a single genus (SI Appendix, Fig. S6).

To understand the relationship between immunogenic outcomes and protein evolution, we developed a maximum-likelihood tree of the cold-shock domain and plotted genera, the percent amino acid similarity of each epitope to the consensus, and immunogenicity by ROS (Fig. 3A). Unlike EF-Tu, CSP domains display intricate clade structure (Fig. 3A). While over 19,000 cold-shock domain-containing proteins were extracted, the 65 csp22 variants tested still managed to capture 75% (14,587/19,423) of the immunogenetic landscape (Fig. 3A-B). Three conserved clades contained divergent CSPs carrying non-immunogenic csp22 epitopes. Non-immunogenic CSPs within Clade 1 were conserved across 62% (781/1261) of *Xanthomonas*, Clade 2 was conserved across 83% (394/475) of *Pectobacterium* and *Erwinia* with a distant relative in *Agrobacterium*, and Clade 3 comprised of 95% (580/612) of actinobacteria (Fig. 3C, right panel).Unlike EF-Tu, almost all genomes carried more than one CSP and of those, 45.7% had at least one non-immunogenic form (Fig. 3C, left panel).

**Fig. 3.**
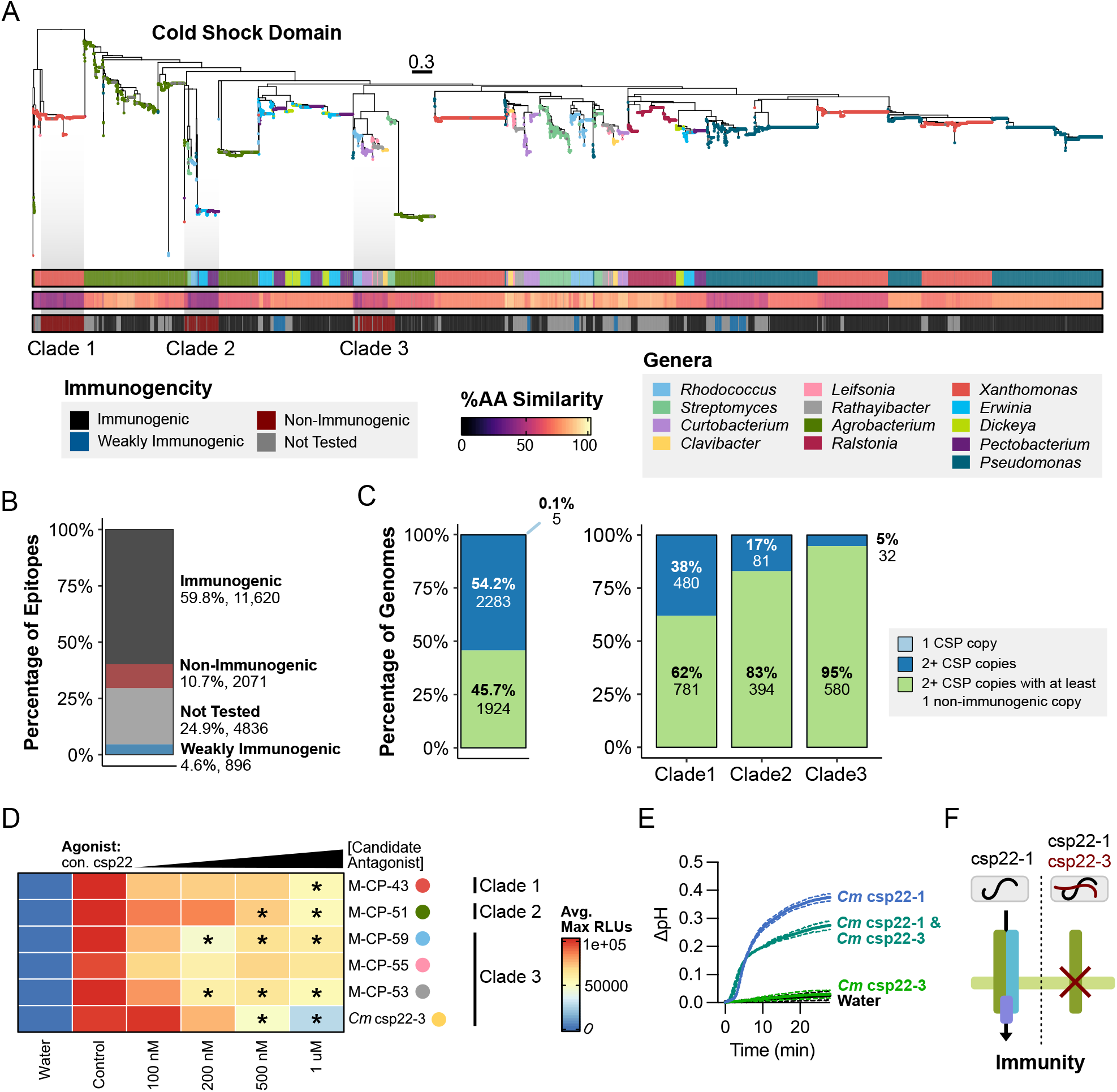
Convergently evolved non-immunogenic CSPs reduce CORE perception. (*A*) Maximum-likelihood phylogenetic tree of CSP domains. Tips were colored by genera, second bar represents percent amino acid similarity to consensus, and third bar indicates immunogenicity based on ROS experiments in Supplemental Figure 6 (200 nM). (*B*) Summary of immunogenic outcomes of csp22 epitopes tested by ROS. (*C*) Summary of immunogenic outcomes in respect to CSP copy number (Left). Summary of immunogenic outcomes from bacteria present across each non-immunogenic Clade in A and their relatives in the same genera (Right). (*D*) ROS screen for immune antagonism in Rio Grande tomato. Leaf disks were incubated overnight with candidate antagonist followed by elicitation with 200 nM agonist peptide. Control denotes positive control (untreated immunogenic agonist) and concentrations listed are of the candidate non-immunogenic antagonist. Maximum RLU averages were adjusted to a scale of 0 to 100,000 based on the controls (water and untreated agonist). Candidate antagonists, labeled by genera, were selected across the three conserved non-immunogenic-derived CSPs from Figure 5. A one-way ANOVA and Tukey’s mean comparison was used to determine significance in respect to the untreated agonist control (denoted *: p < 0.05). (*E*) Alkalization of *Nicotiana tabacum* cv. Bright Yellow (BY-2) suspension cells after MAMP treatment (10 nM *Cm* csp22-1; 10 nM *Cm* csp22-3). Antagonism was assessed by first treating with 50 nM *Cm* csp22-3 for three minutes, then treating with 10 nM agonist *Cm* csp22-1. Two cell aliquots were tested per treatment and the experiment was repeated at least two times. (*F*) Diagram of *Clavibacter* CSP intrabacterial antagonism.

Observing the differences in immunogenic outcomes and sequence diversity, we then assessed conservation and selection of specific residues from the conserved non-immunogenic CSP. Orthologous CSPs were classified using a combination of phylogeny, protein clustering, and motif classification (SI Appendix, Materials and Methods, Fig. S8A-B). We then used an all-by-all BlastP approach to confirm orthology (SI Appendix, Fig. S8C). Next, d_N_/d_S_ was calculated for non-immunogenic and immunogenic CSPs from the same bacterial genera. For non-immunogenic and immunogenic CSP loci, selection was assessed and codon sites that were considered significantly negative or positive were based on a set posterior probability (SI Appendix, Fig. S9A). Among representative immunogenic and non-immunogenic CSP members from the three clades, most sites are either negatively selected (purifying) or neutral with only a very small number of codons displaying positive (diversifying) selection (SI Appendix, Fig. S9A). For all tested immunogenic, weakly-immunogenic, and non-immunogenic epitopes, the RNP-1 motif is highly conserved and under purifying selection (SI Appendix, Fig. S9A-B). Near the N-terminus (highlighted in grey), a subset of residues between positions four and nine may be critical for strong immunity based on the residue changes, R-group chemistry, and immunogenic outcomes (SI Appendix, Fig. S9A). The epitope signature for each clade is unique and derived from distinct paralogs (Fig. 3A, SI Appendix, Fig. S9B).

### Conserved non-immunogenic cps22 variants reduce CORE immune perception

We observed three non-immunogenic CSP clades across different bacterial genera (Fig. 3A, 3C). These non-immunogenic CSPs may be maintained for their role in inhibiting immune perception. To test this, we modified our high-throughput ROS assay to assess epitope antagonism. Briefly, leaf disks were floated on one of two solutions: water or increasing concentrations of the candidate antagonist. After overnight incubation, all liquid was removed, an 100 nM of agonist was used to elicit immunity, and ROS was measured (SI Appendix, Fig. S10A). We selected at least one member from each non-immunogenic clade and used consensus csp22 as the agonist. Most non-immunogenic csp22 epitope variants reduced ROS produced by consensus csp22. The variant conserved in the *Clavibacter* genus, *Cm* csp22-3, displayed the strongest ROS reduction at concentrations which were 2.5x and 5x the respected agonist concentration (Fig. 3D).

*Clavibacter* contains three CSPs with different immunogenic outcomes, immunogenic cspA1 (*Cm* csp22-1), weakly immunogenic cspA2 (*Cm* csp22-2), and non-immunogenic cspB (*Cm* csp22-3) (Fig. 3A, SI Appendix, Fig. S6A). We repeated the antagonism assay with agonist *Cm* csp22-1 alongside two negative controls: elf18 and a scrambled version of *Cm* csp22-3 (designated s-csp22-3) (SI Appendix, Fig. S10). Antagonism was observed via decreased ROS production after incubation with *Cm* csp22-3 but not for the other negative control peptides (SI Appendix, Fig. S10). We additionally tested *Cm* csp22-3 antagonism using BY-2 tobacco cell cultures by measuring alkalization, an output of membrane depolarization through ion fluxes and inhibition of H+-ATPases at the plasma membrane (Fig. 3E) (38,39). As expected, *Cm* csp22-1 (10 nM) was immunogenic and able to induce a pH shift, while *Cm* csp22-3 (10 nM) was not. When 5x *Cm* csp22-3 was added three minutes before *Cm* csp22-1, it was able to decrease the pH shift, showcasing its antagonistic nature (Fig. 3E). Collectively, these data indicate that nonimmunogenic CSP epitope variants can antagonize perception of immunogenic forms encoded in the same genome, which we term intrabacterial antagonism (Fig. 3F).

We also observed that *Streptomyces* carried two copies of EF-Tu with distinct epitope variants (Fig. 2A, SI Appendix, Fig. S12A). One non-immunogenic elf18 variant was conserved across multiple *Streptomyces* genomes (EF-92), with a second non-immunogenic variant found in one genome (EF-97). Using the same ROS antagonism assay, we tested both non-immunogenic epitopes against two different agonists, consensus elf18 and the immunogenic *Streptomyces* variant, EF-77. We tested elf12, a truncated version of consensus elf18, which has been previously reported as a weak antagonist (11). As expected, elf12 was able to weakly reduce ROS induction for either agonist. However, neither EF-92 nor EF-97 displayed consistent ROS reduction at any concentration (SI Appendix, Fig. S12B). Therefore, the capacity for intrabacterial antagonism is MAMP-dependent.

### Intrabacterial antagonism of conserved actinobacterial CSP

Antagonism has been predominantly characterized using peptide assays. Therefore, we investigated intrabacterial antagonism with full-length CSP proteins. The *Clavibacter* genus comprises eight pathogenic species (40,41). Pathogens in the *Clavibacter* genus predominantly colonize the xylem vasculature and upon systemic infection, colonize additional tissues (40). First, conservation and synteny of each *Clavibacter* CSP was analyzed across multiple species. Both immunogenic *cspA1* and non-immunogenic *cspB* were found in all *Clavibacter* genomes, whereas the weakly immunogenic *cspA2* was found in 85% (74/87) (Fig. 4A). Across several species, each gene was highly syntenic (Fig. 4A). Expression of each *CSP* was assessed via qPCR for the tomato pathogen *C. michiganensis* in xylem-mimicking media and compared to a previously published housekeeping gene, *bipA*, in TBY rich medium. While all CSPs are expressed, *cspA2* and *cspB* exhibited higher expression than *cspA1* at both 6 and 24 hours with *cspB* displaying between 2.65 and 3.63x fold change with *cspA1* (Fig. 4B). Furthermore, when recombinant protein CspA1 and CspB were mixed in varying concentrations, we observed a strong reduction in ROS at 2.5x concentrations differences, a realistic difference as expression and protein abundance are correlated in bacteria (Fig. 4C-D) (44).

**Fig. 4.**
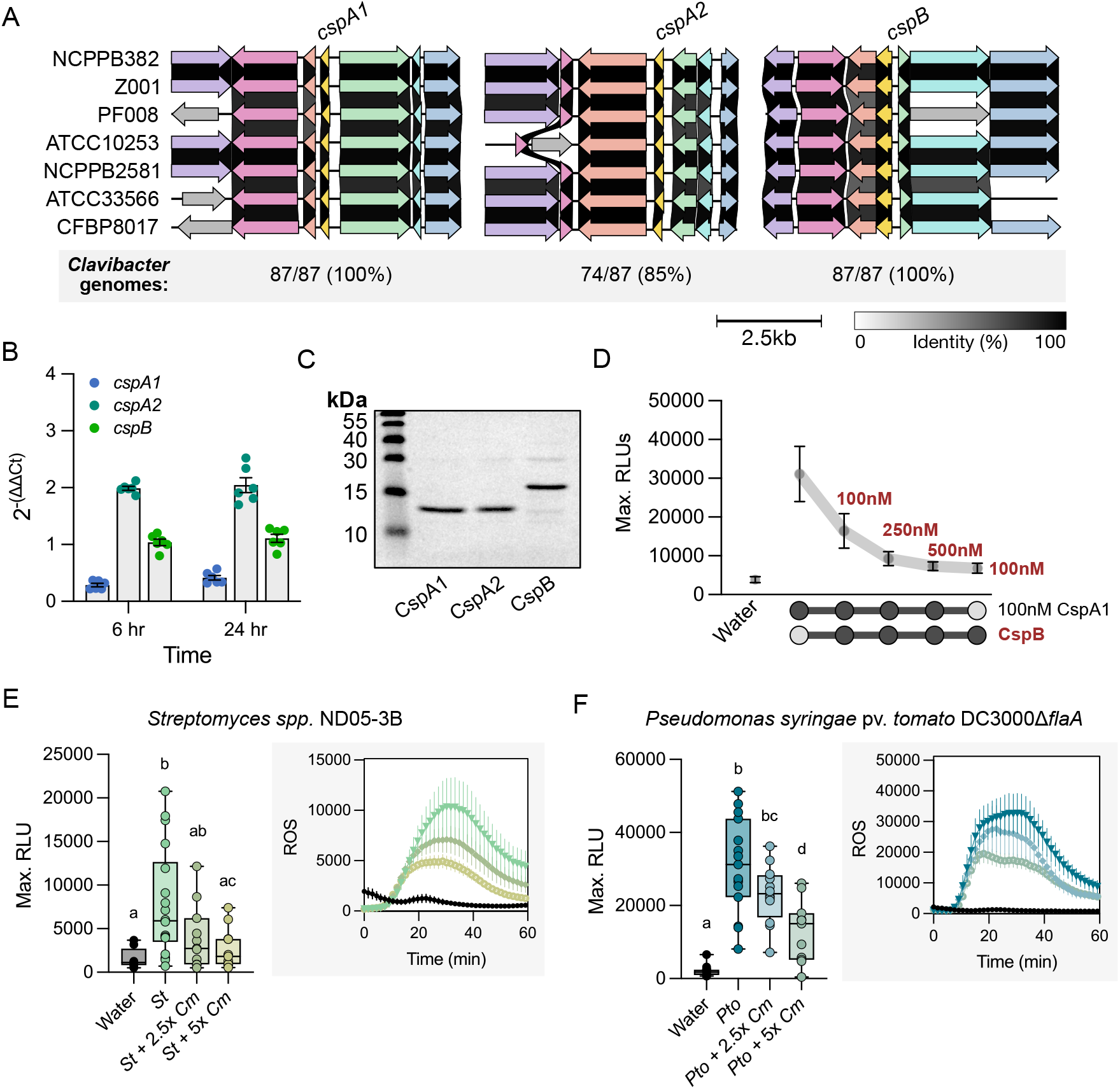
*Clavibacter* CspB is conserved, expressed, and enables immune evasion of CORE. (*A*) Conservation and syntenic gene structure of CSPs in representative *Clavibacter* genomes (*C. michiganensis:* NCPPB382 and Z001; *C. capsici:* PF008; *C. insidiosus:* ATCC10253; *C. nebraskensis:* NCPPB2581; *C. tessellarius:* ATCC33566 and CFBP8017). (*B*) Expression of CSPs from *Clavibacter michiganensis* NCPBB382 in xylem-mimicking media compared to expression of the housekeeping gene, *bipA*, in rich media TBY. Three technical replicates for two independent biological cultures are plotted. The experiment was repeated three times with similar results. Error bars = SEM. (*C*) Coomassie stained SDS-PAGE gel of CSP recombinant proteins purified from *E. coli*. CspA1 is 13 kDa and CspB is 17kDa. (*D*) Maximum ROS production in tomato Rio Grande. Recombinant CSP proteins were mixed in the indicated concentrations at the same time and applied to leaf tissue. Concentrations were 100 nM for *Cm* CspA1 and different concentrations of recombinant non-immunogenic *Cm* CspB (denoted in red font). (*E*) ROS induction of *Streptomyces* sp. ND05-3B (1 ug/mL, *St*) and *C. michiganensis* NCPPB382 bacterial lysates (2.5 and 5 ug/mL, *Cm*) in Rio Grande tomato. (*F*) ROS induction of *Pseudomonas syringae* pv. *tomato* DC3000Δ*flaA* (1 ug/mL) and *C. michiganensis* NCPPB382 bacterial lysates (2.5 and 5 ug/mL) in Rio Grande tomato. Lysates were mixed in the indicated concentrations at the same time and applied to leaf tissue in (*E*) and (*F*). In (*D-F*) the Max. RLUs include an average of four punches per plant, 12 plants per treatment. Error bars = SEM. A one-way ANOVA and Tukey’s mean comparison was used to determine significance, p<0.05.

Next, we wanted to investigate the ability of *C. michiganensis* (*Cm*) cell lysates to suppress *CORE*-dependent immunity from another actinobacteria *Streptomyces spp*. ND05-3B (*St*) as well as the proteobacteria *Pseudomonas syringae* pv. *tomato* DC3000Δ*flaA* (*Pto*). Both *St* and *Pto* carry immunogenic CSPs and lack *cspB* (SI Appendix, Fig. S6A, S11). Both *Clavibacter* and *Streptomyces* are non-flagellated and lack *fliC*; we used a *Pto fliC* deletion mutant (Δ*flaA*). On *Arabidopsis, Cm* lysates strongly induce ROS while *St* and *Pto* weakly induce ROS in an *EFR*-dependent manner (SI Appendix, Fig. S11). *Cm* lysates were unable to induce ROS in tomato cv. Rio Grande or the *core* mutant line (Appendix, Fig. S11). However, *St* and *Pto* were able to induce ROS in a *CORE-*dependent manner. Furthermore, we able to observe reduced ROS production in Rio Grande when *Cm* lysates were mixed with either immunogenic *St* or *Pto* lysates at a 2.5 to 5x concentration (Fig. 4E-F). Collectively, these data demonstrate *Cm* can antagonize *CORE-* mediated perception of *St* and *Pto*.

We were unable to generate a *Clavibacter* knockout of *cspB* using a variety of approaches likely due to the region’s high GC content, ranging from 73 to 78% (42,43). In order to functionally assess intrabacterial CspB antagonism *in planta*, we expressed codon-optimized *cspB* in the tomato pathogen *Pto* DC3000 (Fig. 5). The five CSPs found in the DC3000 genome are immunogenic and expressed *in planta* (Fig. 5A, SI Appendix, Fig. S5A, S11) (44). CSPs are known to act as translational chaperones and antiterminators during environmental conditions such as cold and stress (22). *In vitro* expression of *cspB* was confirmed via western blot (Fig. 5C). *In vitro* growth was not significantly different between *P. syringae* expressing *cspB* or empty vector, indicating that *cspB* expression does not grossly impact fitness (Fig. 5D).

**Fig. 5.**
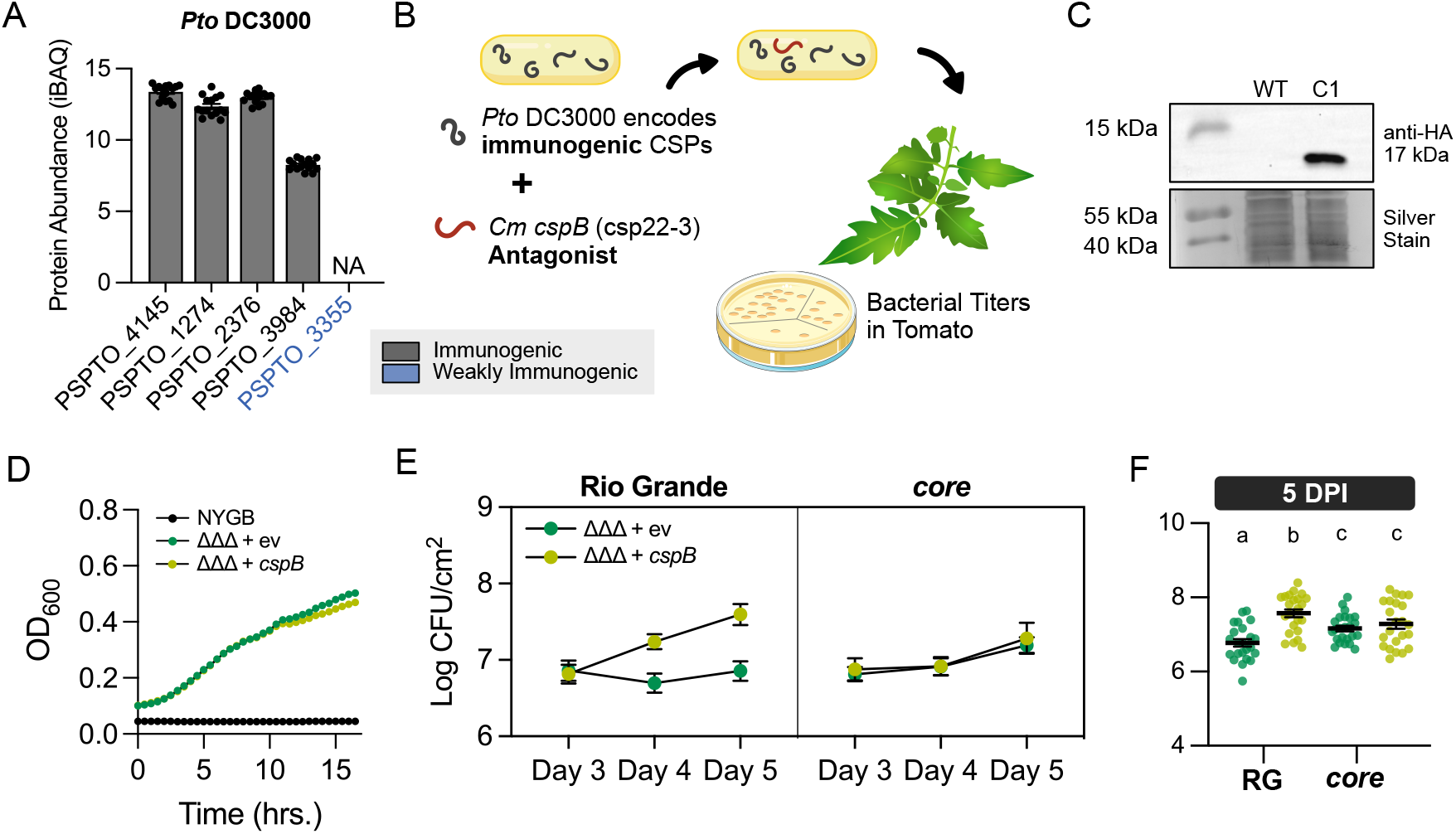
Intrabacterial transfer of CspB antagonizes CORE perception in a Gram-negative tomato pathogen. (*A*) Induction of *Pseudomonas syringae* pv. *tomato* (*Pto*) DC3000 CSPs based on raw count proteome data from *Nobori et al*., 2020.^44^ Variants are labeled by their immunogenicity based on Figure 3 and Supplemental Figure 6. (*B*) Diagram of *Pto* DC3000 transformed with pDSK519 carrying codon-optimized *cspB* where bacterial titers were tested in tomato. (*C*) *In vitro* expression of pDSK519 carrying codon-optimized *cspB* (hereafter pDSK519-*cspB*) in *Pto* DC3000Δ*avrPto*Δ*avrPtoB*Δ*flaA* (ΔΔΔ) (OD_600_ = 1). WT = Wildtype strain DC3000ΔΔΔ. (*D*) Representative plot of DC3000ΔΔΔ *in vitro* growth with pDSK519-ngfp (ev) and with pDSK519-*cspB* in NYGB medium. (*E* and *F*) Bacterial titers of DC3000ΔΔΔ carrying pDSK519-ngfp (ev) or pDSK519-*cspB* in Rio Grande (RG) and *core* mutants three to five days post infiltration. Two leaves were sampled per plant with four plants per strain. The experiment was repeated three times. Error bars = SEM. A one-way ANOVA and Tukey’s mean comparison was used to determine significance, p<0.01.

In order to assess CspB antagonism, we used DC3000Δ*avrPto*Δ*avrPtoB*Δ*flaA* (referred to as DC3000ΔΔΔ) to prevent AvrPto/AvrPtoB effector and flg22 recognition (45). After inoculation with DC3000ΔΔΔ, both *core* mutant lines exhibited increased disease symptoms and bacterial titers compared to the wild-type Rio Grande control (SI Appendix, Fig. S5C-D). These data demonstrate that tomato *CORE* impacts *P. syringae* colonization. Next, we characterized bacterial titer development between three- and five-days post infiltration for DC3000ΔΔΔ expressing *cspB*. In Rio Grande, DC3000ΔΔΔ expressing *cspB* exhibited higher bacterial titers at four- and five-days post-inoculation compared to DC3000ΔΔΔ carrying empty vector (Fig. 5E). It is likely we detected a late increase in bacterial titer because cells require lysis to release CSPs and a minimum concentration of CspB is required to overcome immunogenic CSPs from *P. syringae*. Importantly, when either strain was inoculated on the *core* mutant line, there were no significant differences in bacterial titers between DC3000ΔΔΔ expressing *cspB* and DC3000ΔΔΔ carrying empty vector over the course of five days (Fig. 5F). Thus, intrabacterial expression of CspB is sufficient to inhibit CORE perception. Collectively, these data demonstrate that *Clavibacter* CspB is an intrabacterial antagonist.

## Discussion

Natural epitope variation showcases outcomes which can serve to inform evolution, mechanistic interactions, and improve engineering approaches. We surveyed thousands of plant-associated bacteria and found variability in epitope sequence and CNV in a MAMP- and genera-dependent manner. Since natural variation was relatively low, characterizing the immune landscape for MAMP perception across thousands of bacteria is possible. Our work showcases the predictable nature of epitope evolution, enabling future rapid prediction of MAMP immunogenic outcomes. We uncovered a new mechanism of immune evasion, intrabacterial antagonism, which demonstrates that all genome-encoded MAMPs need to be considered when characterizing bacterial-plant outcomes.

Millions of years of evolution led to bacterial divergence into different phyla. Recent work has assessed sequence variation for the flagellin epitope, flg22, revealing diverse yet constrained variation (32,35). Colaianni et al. identified 268 unique variants from 627 *Arabidopsis*-associated genomes. We also identified a similar number of flg22 epitope variants (254) from 4,228 genomes, highlighting the constrained nature of epitope variation across plant-associated bacteria (35). In contrast, 1,059 flg22 variants were identified from 1,414 genomes from representative proteobacterial genomes across diverse lifestyles (32). These studies revealed differential host immunogenic outcomes based on specific residue positions. In flg22, sites 14 and 15 can have a direct impact on immune perception but consequently impair motility (8,32,46). Therefore, the prevalence of some mutations may be constrained due to negative fitness effects.

Epitope diversification may shift when multiple copies are present (35,47). Gene expansion can impact both protein function and epitope variation. For some *Pseudomonas* strains, FliC-1 is critical for both plant immune perception and motility, while FliC-2 is poorly expressed and only weakly induces immune perception (47). For FliC-2, EF-Tu and CSPs, certain residues changes in their respective epitope sequences were specific to additional copies (Fig. 2, 3) (47). With the exception of *Streptomyces*, most other bacterial genera carry a single EF-Tu locus and polymorphisms were highly constrained and predictable in position, likely reflective of the protein’s critical function to bacterial life (SI Appendix, Fig. S4A) (48). In contrast, CSPs are predominantly multi-copy genes, which likely influences their expanded variation (Fig. 1, 3). Individual CSPs can have minimal effects on bacterial survival when knocked out, although they can affect pathogen virulence (22,27,28,49).

Some MAMP-encoded genes have multiple epitopes which can be recognized by different receptors (50-52). Different epitopes of FliC can be detected from distinct PRRs in *Arabidopsis*, tomato, rice, and vertebrates (15,21,53). Each plant genome contains hundreds of candidate PRRs; thus, it is possible that linage specific, convergent evolved receptors is common and the broad conservation of FLS2 is the exception (54). While *CORE* is restricted to the Solanaceae, other genotypes in the Vitaceae and Rutaceae families respond to csp22 variants (10,55). Considering CSP diversity, it is likely other convergently evolved PRRs recognize and respond to csp22 variants.

MAMP-derived genes have been rarely assessed beyond the first copy. One study assessed how a small number of *Pseudomonas* strains carrying two *FliC* genes exhibited differential immune outputs when individually assessed (47). For bacteria carrying one *FliC*, antagonism has been demonstrated against other genera (35,47). Here, we discovered actinobacteria contain multiple encoded CSPs where one copy blocks perception of additional immunogenic forms, thus functioning as intrabacterial antagonist (Fig. 3–5). Gram-positive pathogens lack the type three secretion system and cannot deliver immune suppressing effectors directly into host cells (40). Therefore, intrabacterial CSP antagonism provides an additional mechanism of immune suppression. Intrabacterial MAMP antagonism may also influence community dynamics or mixed infections.

While total variation differs between MAMPs, csp22 epitopes exhibited the most variation (SI Appendix, Fig. S2). MAMP epitopes are proposed to undergo an arms race and are presumed to be under diversifying selection (56,57). However, EF-Tu is under purifying selection (57). Furthermore, across multiple CSP orthologs present in genera with both immunogenic and non-immunogenic forms, most codons displayed either neutral or purifying selection (SI Appendix, Fig. S9A). There are many CSP copies per genome with potentially different functions, frequently ranging from three to nine, thus the selection pressure on any one CSP may be less compared to other loci (Fig. 3A, SI Appendix, Fig. S2). Additionally, perception of CSPs is thought to require cell lysis for perception, which may impact diversifying selection.

By characterizing the consequence of natural epitope variation on immune perception, alternate immune outcomes have been revealed. While reduction in ROS production by some epitope variants was measurable, we failed to characterize any elf18 or csp22 epitopes which displayed statistically significantly higher ROS production compared to the consensus controls (SI Appendix, Fig. S4A, Fig. S6A). This may reveal that a minimal number of residues at specific positions are sufficient for strong complex formation and immune induction. Some flg22 variants, called deviant peptides, can uncouple immune outputs; they are able to induce early-stage ROS production but fail to induce late-stage immune responses such as SGI (35). This phenomenon of uncoupled immune outputs was also found for some elf18 variants (Fig. 2). Two weakly immunogenic variants by ROS, one from *Xanthomonas* and one from *Streptomyces*, were unable to induce SGI, callose deposition, or MAPK phosphorylation. It is likely MAMPs beyond flg22 and elf18 may also act as deviants, representing another strategy to reduce strong immune outputs. We envision two possible explanations for deviant peptide evolution. One, we may be capturing a snapshot of peptide evolution toward either maskers or antagonists. Two, these deviant peptides may occupy the space of the receptor complex, limiting robust immune outputs by other immunogenic peptides. These strategies are not unique to pathogens alone; beneficial microbial communities have been shown to be enriched in antagonistic FliC variants, which block perception of FLS2 (35).

A common strategy to confer disease resistance is to transfer receptors between plant genotypes. *EFR* has been transferred to tomato, citrus, wheat, and apple (58-61). In each case, *EFR* transgenic plants are able to significantly restrict pathogen titers and disease progression. This conclusion is congruent with our elf18 screen since most EF-Tu epitopes (92%) induce strong immune responses in *A. thaliana* (Fig. 2). In contrast, prevalent non-immunogenic epitopes exist from FliC and CSPs, of which some block receptor recognition (Fig. 3, SI Appendix, Fig. S2) (30,32,35,62). Therefore, careful consideration of pathogen epitope variation should be considered for receptor transfer and subsequent engineering (63). We encourage future work using this dataset to inform which receptors may be optimal for pathogen control. In concert, large epitope variant databases may enable new synthetic receptor engineering via protein modeling of receptor structure. Rational receptor design focused on contact with low-polymorphic ligand residues may delay evolution of pathogen evasion.

## Materials and Methods

Bacterial genomes were pulled from NCBI’s Refseq and bacterial epitopes were mined via BlastP, local alignment, and custom R scripts for filtering (Fig. 1A, Dataset S1) (6,11,13,16,17). Phylogenetic trees for bacteria relatedness were built using GToTree and protein trees were built using MAFFT for sequence alignment and IQ-TREE tree building. Epitopes of interests that were assessed *in planta* were chosen based on abundance, epitope sequence, and gene annotation. ROS, MAPK, seedling growth inhibition, callose deposition, and alkalinization were measured similarly as previously described with some minor modifications (13, 31, 35). Gene structure was assessed in *C. michiganensis* using BlastN and Clinker. Measurement of *C. michiganensis* CSP expression was conducted similarly as previously described (64). Collection of bacterial lysates were cultured in liquid media or on plates, collected, and processed in lysis buffer via a Emusiveflex-C3 High-Pressure Homogenizer. Recombinant protein was expressed in *E. coli* and collected using standard techniques (7). Bacterial titers of *P. syringae* in tomato was conducted similarly as described before (14). Detailed *Materials and Methods* are found in SI Appendix.

## Supporting information

Dataset S01, genome accessions

## Acknowledgments

This research was supported by NIH 1R35GM136402, US-Israel BARD IS-5499-22 to G.C. and USDA-NIFA 2021-67034-35049, UC Davis Henry A. Jastro Scholarship, and UA Local 290 Scholarship to D.M.S. A.M.P. was supported by Fundación Alfonso Martín Escudero Postdoctoral fellowship. A.J.W. was supported by startup funding from the Department of Botany and Plant Pathology at Oregon State University. C.R. was supported by UC Davis Dean’s Distinguished Graduate Fellowship. We thank Marta Bjornson and Jeff Dangl for their thoughtful discussions of this work and Emily Fucarino for plant growth. Jianfeng Xu (Arkansas Biosciences Institute) provided tobacco BY-2 callus, Imran Khan and Karen MacDonald (UC Davis) and Fumi Fukada (Okayama University) provided guidance with maintaining cell cultures, Jeff Chang (Oregon State University) provided the helper plasmid strains pRK2013 and pRK600, Anjali Iyer-Pascuzzi (Purdue University) provided the flgII-28 peptide, and Christopher Clarke (USDA Agricultural Research Service, Beltsville, MD) for the *Streptomyces* strain ND05-3B. We thank Nitzan Shabek and Shahid Siddique (UC Davis) for the usage of the EmusiveFlex and Biotek plate reader, respectively.

## Data, Materials, and Software Availability

Accessions of genomes used in this study can be found Dataset S1. Plasmids pDSK519-cspB (#207162), pRSET-Cm-CspA1 (#215398), and pRSET-Cm-CspB (#215400) can be found in Addgene. All raw data can be found in Zenodo (Doi: 10.5281/zenodo.10724865). Complete details and code can be found in the following GitHub repository: DanielleMStevens/Mining_Known_MAMPs. An HTML file with all MAMPs mined can be found in Zenodo (Doi: 10.5281/zenodo.10724865).

## References

1. D. C. Lewis, D. M. Stevens, H. Little, G. L. Coaker, R. M. Bostock, Overlapping Local and Systemic Defense Induced by an Oomycete Fatty Acid MAMP and Brown Seaweed Extract in Tomato. Mol. Plant-Microbe Interact. 36, 359–371 (2023).

2. B. P. M. Ngou, P. Ding, J. D. G. Jones, Thirty years of resistance: Zig-zag through the plant immune system. The Plant Cell 34, 1447–1478 (2022).

3. D. Ge, I.-C. Yeo, L. Shan, Knowing me, knowing you: Self and non-self recognition in plant immunity. Essays Biochem 66, 447–458 (2022).

4. S. Lolle, D. Stevens, G. Coaker, Plant NLR-triggered immunity: from receptor activation to downstream signaling. Curr Opin Immunol 62, 99–105 (2020).

5. X. Yu, B. Feng, P. He, L. Shan, From Chaos to Harmony: Responses and Signaling Upon Microbial Pattern Recognition. Annu Rev Phytopathol 55, 1–29 (2016).

6. G. Felix, J. D. Duran, S. Volko, T. Boller, Plants have a sensitive perception system for the most conserved domain of bacterial flagellin. Plant J 18, 265–276 (1999).

7. Y. Wei, et al., An immune receptor complex evolved in soybean to perceive a polymorphic bacterial flagellin. Nat Commun 11, 3763 (2020).

8. K. Parys, et al., Signatures of antagonistic pleiotropy in a bacterial flagellin epitope. Cell Host Microbe 29, 620-634.e9 (2021).

9. L. Wu, H. Xiao, L. Zhao, Q. Cheng, CRISPR/Cas9-mediated generation of fls2 mutant in Nicotiana benthamiana for investigating the flagellin recognition spectrum of diverse FLS2 receptors. Plant Biotechnol J20, 1853–1855 (2022).

10. J. Trinh, et al., Variation in microbial feature perception in the Rutaceae family with immune receptor conservation in citrus. Plant Physiol. (2023) doi:10.1093/plphys/kiad263.

11. G. Kunze, et al., The N Terminus of Bacterial Elongation Factor Tu Elicits Innate Immunity in Arabidopsis Plants. Plant Cell Online 16, 3496–3507 (2004).

12. C. Zipfel, et al., Perception of the Bacterial PAMP EF-Tu by the Receptor EFR Restricts Agrobacterium-Mediated Transformation. Cell 125, 749–760 (2006).

13. G. Felix, T. Boller, Molecular Sensing of Bacteria in Plants THE HIGHLY CONSERVED RNA-BINDING MOTIF RNP-1 OF BACTERIAL COLD SHOCK PROTEINS IS RECOGNIZED AS AN ELICITOR SIGNAL IN TOBACCO. J Biol Chem 278, 6201–6208 (2003).

14. L. Wang, et al., The pattern-recognition receptor CORE of Solanaceae detects bacterial cold-shock protein. Nat Plants 2, 16185 (2016).

15. S. R. Hind, et al., Tomato receptor FLAGELLIN-SENSING 3 binds flgII-28 and activates the plant immune system. Nat. Plants 2, 16128 (2016).

16. R. Cai, et al., The Plant Pathogen Pseudomonas syringae pv. tomato Is Genetically Monomorphic and under Strong Selection to Evade Tomato Immunity. PLoS Pathog. 7, e1002130 (2011).

17. H. Böhm, et al., A Conserved Peptide Pattern from a Widespread Microbial Virulence Factor Triggers Pattern-Induced Immunity in Arabidopsis. Plos Pathog 10, e1004491 (2014).

18. I. Albert, et al., An RLP23–SOBIR1–BAK1 complex mediates NLP-triggered immunity. Nat Plants 1, nplants2015140 (2015).

19. M. F. Seidl, G. V. den. Ackerveken, Activity and Phylogenetics of the Broadly Occurring Family of Microbial Nep1-Like Proteins. Annu Rev Phytopathol 57, 367–386 (2019).

20. F. F. V. Chevance, K. T. Hughes, Coordinating assembly of a bacterial macromolecular machine. Nat. Rev. Microbiol. 6, 455–465 (2008).

21. Y. Rossez, E. B. Wolfson, A. Holmes, D. L. Gally, N. J. Holden, Bacterial Flagella: Twist and Stick, or Dodge across the Kingdoms. PLoS Pathog. 11, e1004483 (2015).

22. Y. Zhang, C. A. Gross, Cold Shock Response in Bacteria. Annu. Rev. Genet. 55, 1–24 (2021).

23. B. Schmid, et al., Role of Cold Shock Proteins in Growth of Listeria monocytogenes under Cold and Osmotic Stress Conditions. Appl. Environ. Microbiol. 75, 1621–1627 (2009).

24. L. P. Burbank, D. C. Stenger, A Temperature-Independent Cold-Shock Protein Homolog Acts as a Virulence Factor in Xylella fastidiosa. Mol. Plant-Microbe Interact. 29, 335–344 (2016).

25. A. K. Eshwar, C. Guldimann, A. Oevermann, T. Tasara, Cold-Shock Domain Family Proteins (Csps) Are Involved in Regulation of Virulence, Cellular Aggregation, and Flagella-Based Motility in Listeria monocytogenes. Front. Cell. Infect. Microbiol. 7, 453 (2017).

26. A. Catalan-Moreno, et al., One evolutionarily selected amino acid variation is sufficient to provide functional specificity in the cold shock protein paralogs of Staphylococcus aureus. Mol. Microbiol. 113, 826–840 (2020).

27. W. Wei, T. Sawyer, L. Burbank, Csp1, a Cold Shock Protein Homolog in Xylella fastidiosa Influences Cell Attachment, Pili Formation, and Gene Expression. Microbiol. Spectr. 9, e01591–21 (2021).

28. L. Wu, L. Ma, X. Li, Z. Huang, X. Gao, Contribution of the cold shock protein CspA to virulence in Xanthomonas oryzae pv. oryzae. Mol. Plant Pathol. 20, 382–391 (2019).

29. K. L. Harvey, V. M. Jarocki, I. G. Charles, S. P. Djordjevic, The Diverse Functional Roles of Elongation Factor Tu (EF-Tu) in Microbial Pathogenesis. Front Microbiol 10, 2351 (2019).

30. S. Wang, et al., Rice OsFLS2-Mediated Perception of Bacterial Flagellins Is Evaded by Xanthomonas oryzae pvs. oryzae and oryzicola. Mol. Plant 8, 1024–1037 (2015).

31. Y. Wei, et al., The Ralstonia solanacearum csp22 peptide, but not flagellin-derived peptides, is perceived by plants from the Solanaceae family. Plant Biotechnol J 16, 1349–1362 (2018).

32. J. H. T. Cheng, M. Bredow, J. Monaghan, G. C. diCenzo, Proteobacteria Contain Diverse flg22 Epitopes That Elicit Varying Immune Responses in Arabidopsis thaliana. Mol Plant-microbe Interactions 34, 504–510 (2021).

33. U. Fürst, et al., Perception of Agrobacterium tumefaciens flagellin by FLS2XL confers resistance to crown gall disease. Nat. Plants 6, 22–27 (2020).

34. P. Buscaill, R. A. L. van der. Hoorn, Defeated by the nines: nine extracellular strategies to avoid microbe-associated molecular patterns recognition in plants. Plant Cell 33, koab109 (2021).

35. N. R. Colaianni, et al., A complex immune response to flagellin epitope variation in commensal communities. Cell Host Microbe 29, 635-649.e9 (2021).

36. B. Castro, et al., Stress-induced reactive oxygen species compartmentalization, perception and signalling. Nat. Plants 7, 403–412 (2021).

37. F. Yang, G. Li, G. Felix, M. Albert, M. Guo, Engineered Agrobacterium improves transformation by mitigating plant immunity detection. New Phytol (2022) doi:10.1111/nph.18694.

38. M.M. Atkinson, J.-S. Huang, and K.A. Knopp, The Hypersensitive reaction of tobacco to Pseudomonas syringae pv. Pisi: Activation of a plasmalemma K+/H+ Exchange Mechanism. Plant Physiol. 79, 843–847 (1985).

39. J. M. Elmore, G. Coaker, The Role of the Plasma Membrane H+-ATPase in Plant–Microbe Interactions. Mol Plant 4, 416–427 (2011).

40. S. P. Thapa, et al., The Evolution, Ecology, and Mechanisms of Infection by Gram-Positive, Plant-Associated Bacteria. Annu Rev Phytopathol 57, 341–365 (2019).

41. D. Arizala, S. Dobhal, A. M. Alvarez, M. Arif, Elevation of Clavibacter michiganensis subsp. californiensis to species level as Clavibacter californiensis sp. nov., merging and re-classification of Clavibacter michiganensis subsp. chilensis and Clavibacter michiganensis subsp. phaseoli as Clavibacter phaseoli sp. nov. based on complete genome in silico analyses. Int. J. Syst. Evol. Microbiol. 72, (2022).

42. L. Chalupowicz, et al., Differential contribution of Clavibacter michiganensis ssp. michiganensis virulence factors to systemic and local infection in tomato. Mol Plant Pathol 18, 336–346 (2017).

43. D. M. Stevens, A. Tang, G. Coaker, A Genetic Toolkit for Investigating Clavibacter Species: Markerless Deletion, Permissive Site Identification, and an Integrative Plasmid. Mol Plant-microbe Interactions 34, 1336–1345 (2021).

44. T. Nobori, et al., Multidimensional gene regulatory landscape of a bacterial pathogen in plants. Nat. Plants 6, 883–896 (2020).

45. B. H. Kvitko, et al., Deletions in the Repertoire of Pseudomonas syringae pv. tomato DC3000 Type III Secretion Effector Genes Reveal Functional Overlap among Effectors. Plos Pathog 5, e1000388 (2009).

46. K. Naito, et al., Amino Acid Sequence of Bacterial Microbe-Associated Molecular Pattern flg22 Is Required for Virulence. Mol. Plant-Microbe Interact. 21, 1165–1174 (2008).

47. Y. Luo, J. Wang, Y.-L. Gu, L.-Q. Zhang, H.-L. Wei, Duplicated Flagellins in Pseudomonas Divergently Contribute to Motility and Plant Immune Elicitation. Microbiol. Spectr. 11, e03621–22 (2023).

48. P. J. P. Teixeira, N. R. Colaianni, C. R. Fitzpatrick, J. L. Dangl, Beyond pathogens: microbiota interactions with the plant immune system. Current Opinion in Microbiology 49, (2019).

49. Y. Liu, et al., The cold shock family gene cspD3 is involved in the pathogenicity of Ralstonia solanacearum CQPS-1 to tobacco. Microb. Pathog. 142, 104091 (2020).

50. T. Furukawa, H. Inagaki, R. Takai, H. Hirai, F.-S. Che, Two Distinct EF-Tu Epitopes Induce Immune Responses in Rice and Arabidopsis. Mol Plant-microbe Interactions27, 113–124 (2014).

51. Y. Katsuragi, et al., CD2-1, the C-Terminal Region of Flagellin, Modulates the Induction of Immune Responses in Rice. Mol Plant-microbe Interactions 28, 648–658 (2015).

52. M. de O. Andrade, et al., Suppression of citrus canker disease mediated by flagellin perception. Mol. Plant Pathol.24, 331–345 (2023).

53. J. Fliegmann, G. Felix, Immunity: Flagellin seen from all sides. Nat. Plants 2, 16136 (2016).

54. B. P. M. Ngou, R. Heal, M. Wyler, M. W. Schmid, J. D. G. Jones, Concerted expansion and contraction of immune receptor gene repertoires in plant genomes. Nat. plants 8, 1146–1152 (2022).

55. L. P. Burbank, J. Ochoa, Evidence for Elicitation of an Oxidative Burst in Vitis vinifera by Xylella fastidiosa Cold Shock Protein Peptide csp20. PhytoFrontiers 2, 339–341 (2022).

56. H. C. McCann, H. Nahal, S. Thakur, D. S. Guttman, Identification of innate immunity elicitors using molecular signatures of natural selection. Proc National Acad Sci 109, 4215–4220 (2012).

57. N. Eckshtain-Levi, A. J. Weisberg, B. A. Vinatzer, The population genetic test Tajima’s D identifies genes encoding pathogen-associated molecular patterns and other virulence-related genes in Ralstonia solanacearum. Molecular Plant Pathology 19, (2018).

58. S. Lacombe, et al., Interfamily transfer of a plant pattern-recognition receptor confers broad-spectrum bacterial resistance. Nat Biotechnol 28, 365 (2010).

59. H. Schoonbeek, et al., Arabidopsis EF-Tu receptor enhances bacterial disease resistance in transgenic wheat. New Phytol 206, 606–613 (2015).

60. L. K. Mitre, et al., The Arabidopsis immune receptor EFR increases resistance to the bacterial pathogens Xanthomonas and Xylella in transgenic sweet orange. Plant Biotechnol J 19, 1294– 1296 (2021).

61. S. Piazza, et al., The Arabidopsis pattern recognition receptor EFR enhances fire blight resistance in apple. Hortic Res 8, 204 (2021).

62. C. Pfund, et al., Flagellin Is Not a Major Defense Elicitor in Ralstonia solanacearum Cells or Extracts Applied to Arabidopsis thaliana. Mol. Plant-Microbe Interact. 17, 696–706 (2004).

63. A. Schultink, A. D. Steinbrenner, A playbook for developing disease-resistant crops through immune receptor identification and transfer. Curr. Opin. Plant Biol.62, 102089 (2021).

64. N. Jiang, et al., Evaluation of suitable reference genes for normalization of quantitative reverse transcription PCR analyses in Clavibacter michiganensis. Microbiology open 8, e928 (2019).

