## Supplementary materials and methods, Fig S1-S12, Table S1-S3 for "Natural variation of immune epitopes reveals intrabacterial antagonism"

### Supporting Information for
Natural variation of immune epitopes reveals intrabacterial antagonism.

Danielle M. Stevens, Alba Moreno-Pérez, Alexandra J. Weisberg, Charis Ramsing, Judith Fliegmann, Ning Zhang, Melanie Madrigal, Gregory Martin, Adam Steinbrenner, Georg Felix, and Gitta Coaker

**This PDF file includes:**

SI Materials and Methods
Figures S1 to S12
Tables S1 to S3
SI References

**Other supporting materials for this manuscript include the following:**

Datasets S1

#### SI Materials and Methods

##### *Collection of Genomes and MAMPs*

Bacterial genomes were pulled from NCBI's RefSeq data repository. Briefly, using the –dry-run command for the ncbi-genome-download package (v0.3.0), candidate genomes were queried and manually assessed for associations with plants and/or agriculture. Once a list for each major genre in this study, *Clavibacter, Leifsonia, Curtobacterium, Streptomyces, Rathayibacter, Rhodococcus, Agrobacterium, Ralstonia, Xanthomonas, Pseudomonas, Pectobacterium, Dickeya, and Erwinia,* was collected, all Genbank, protein fasta and whole genome fasta files were downloaded for each Refseq accession number (Dataset S1) using ncbi-genome-download. Consensus MAMPs csp22 (AVGTVKWFNAEKGFGFITPDDG), elf18 (SKEKFERTKPHVNVGTIG), flg22 (QRLSTGSRINSAKDDAAGLQIA), flgII-28 (ESTNILQRMRELAVQSRNDSNSATDREA), and nlp20 (AIMYSWYFPKDSPVTGLGHR) were used to build a custom blast database via makeblastdb. Each protein fasta file was queried against the database using Blastp with the following modifications: -task blastp-short -xdrop_gap_final 1000 -soft_masking false -evalue 1e-4 (1). Secondly, MAMPs were found by local alignment to proteins with typical gene annotations (flagellin, cold shock protein, elongation factor, and necrosis-and-ethylene inducing peptide). Custom R scripts were built process blast results, filter for missing MAMP hits based on protein annotation, correction for gaps on polymorphic ends, filtering for off-targets based on a signature motif for each MAMP and removing hits from partial protein annotations (Fig. 1A). Motifs used to filter for off-targets include: csp22: "KGFGF", elf18: "NXGTXG", flg22: "SXGXXXXXXXXXAA", and flgII-28: "LQRXRXL". Table S1 includes the number of MAMPs detected using this pipeline.

##### *Comparing genome similarity and MAMP hits to filter for clonality*

To assess intra-genera genome diversity, average nucleotide identity via fastANI (v1.32) was calculated in an all-by-all manner within each genus using default parameters (Extended Data Fig. 1; Doi: 10.5281/zenodo.10724865) (2,3). Custom R scripts were used to parse ANI values. Genomes which shared an ANI value of 99.999 percent or higher and shared identical MAMP sequences were considered clonal. Of those that were categorized as clonal, only one representative genome was used. Full description of the methods of ANI analysis can be found in the Github, DanielleMStevens/Mining_Known_MAMPs, particularly under methods and R scripts #10 through 13. Genome accessions which were found to be clonal and removed from subsequent analysis can be found in Supplemental Table 1. The final MAMP list was outputted into an html file using R package DT (v0.20) and can be found in Zenodo (Doi: 10.5281/zenodo.10724865) (3).

##### *Determining epitope diversity*

After all epitopes were extracted, we assessed epitope diversity across a variety of scales. First, we determined how similar the epitopes are in comparison to the consensus peptide were by calculating protein similarity via BlastP as listed in Collection of Genomes and MAMPs section or by local-alignment and calculated via R package Biostrings (v2.58.0), which was used when the BlastP results dropped ends which were polymorphic in sequence compared to the consensus query used (R script #18). Two, to understand epitope variation which did not rely on consensus, a custom R script (#17) was developed to determine all the unique combinations found and performed all-by-all local alignment and amino acid similarity calculations. All-by-all similarity values were plotted as a heatmap using ComplexHeatmap (v2.5.1) and circlize (v0.4.8) (4,5). Three, epitope weblogos were made using ggseqlogo (v0.1) (R script #26) (6). Four, the number of epitope variants (described combination) was determined using a custom script (R script #19) and plotted using ggplot and ggbreak (v0.1.1) (7).

##### *Epitope Selection Criteria*

To determine which epitopes to evaluate, we first selected epitope variants based on their abundance in the dataset (those which occurred at least 10 to 100x). Additionally, epitope variants which were encoded in additional paralogs with unique evolutionary history, domain structure, and/or annotation were selected. In total, we selected 25 elf18 variants and 65 csp22 variants for synthesis and subsequent testing in plants (Table S2).

***Construction of Phylogenetic Trees***

To assess the relatedness of plant-associated bacteria in respect to their MAMP abundance, a phylogenomic tree was constructed using GToTree (v1.6.31) with the MAMPs mined plotted (8). GToTree is an automated workflow for developing phylogenetic trees based on a pre-set list of validated conserved genes (74 for bacteria). First, a list of paths of all GenBank files was outputted into a text file. Genomes which were indicated to be clonal by ANI and MAMPs detected were manually removed. GToTree was ran on the text file with the GenBank file paths using default parameters with the following modifications: -H Bacteria -G 0.2 -k.

Phylogenetic trees were built from the CSP domain and full-length sequences of EF-Tu. Proteins sequences were collected into a fasta file, and a multiple sequence alignment was built via MAFFT (v7.310) using the following parameters: --reorder --thread 12 --maxiterate 1000 –localpair (9). For CSP phylogeny, some CSPs that were longer than 180 amino acids in length were discovered to have additional domains. Therefore, the conserved cold shock protein domain was extracted across all CSPs using via HMMER (v3.3.2) using hmmsearch with the following parameters, -E 1 --domE 1 --incE 0.01 --incdomE 0.04, and the hmm model specific for cold shock domain (3,10). These hits were reformatted using the esl-reformat command in HMMER under default settings before a multiple sequence alignment was build using MAFFT as described above. For the EF-Tu phylogeny, no additional modifications before multiple sequence alignment were necessary. Maximum-likelihood phylogenetic trees were built from the multiple sequence alignments without trimming gaps using IQ-TREE2 (v2.1.2) with the following parameters: -st AA -bb 1000 -mtree -nt 12 -safe (11).

***Classification of cold shock proteins and subsequent selection tests***

To assess selection of orthologous CSP, all loci were grouped into equivalent types. First, all epitope containing hits with annotations including the key words ‘cold, shock, and/or antiterminator’ were filtered into a fasta file in a genera dependent manner using a custom R script (R script #21, 23). The collected CSPs for each genus were that ran through three pipelines. One, to determine the number of CSP clusters, full length sequences were run through mmseq2 (v13.45111) using the easy-cluster command withing the following parameters: clusterRes tmp –min-seq-id 0.5 -c 0.8 –cov-mode 1 (12). Two, full length sequences were passed through MEME Suite Motif Discovery (v5.5.1) under the following parameters: Classic mode, any number of repetitions (anr), and the number of motifs selected based on the number of clusters outputted by mmseq2 and manual inspection (13). The XML output was downloaded for each genera’s CSP loci and processed via the R package ggmotif (v0.2.1) (14,15). Third, the CSP domain was extracted via HMMER (v3.3.2) using hmmsearch with the following parameters, -E 1 --domE 1 --incE 0.01 --incdomE 0.04, and the hmm model specific for cold shock domain can be download (3,10). The CSP domains from each genus were aligned using MAFFT using –auto and a phylogenetic tree was built using FastTree2 (9,16). Information extracted from MEME and mmseq2 analysis was plotted onto the phylogenetic tree to cross reference the number of CSP types identified (R script #24). The R packages phangorn (v2.7.1), treeio (v1.14.4), ggtree (v3.1.2.991), ggnewscale (v0.4.5), and ggtreeExtra (v1.0.4) were used to parse and plot the treefile or newick file (17-20). Details on parameters for plotting can be found in the custom R script #14 and 22.

To confirm within-genera grouped CSPs were the same ortholog, an all-by-all BlastP search was performed using the orthologr R package (v0.4.0) (R script #25) (21). The coding sequences of associated orthologs CSP within a genus were extracted for selection analyses via dn/ds ratio. MACSE (v. 2.07) with the parameters “-prog alignSequences” was used to generate codon alignments from nucleotide gene sequences (22). IQ-TREE (v.1.6.12) with the default parameters was used to generate phylogenies for each dataset (23). HYPHY (v.2.5.48) with the parameter “cln” was used to remove stop codons from alignments (24). AliView (v.1.26) was used to manually inspect alignments and remove gaps present in >75% of sequences (25). The HYPHY algorithm FUBAR, with the codon alignment and gene phylogeny as input and a posterior probability threshold of 0.9, was used to identify sites under purifying or positive selection (24).

***Plant growth conditions***

*Arabidopsis* Col-0 and *efr*-1 transgenics were grown in Sunshine Mix soil in chambers at 10/12-h light/dark cycles, 23°C, about 100 μmol/m$^2$/s light intensity, and at 70% humidity. For reactive oxygen species (ROS) assays, plants were grown to four to five-weeks (26). For MAPK induction, callose deposition and seedling growth inhibition assays, *Arabidopsis thaliana* Col-0 and *efr* seeds were surface sterilized with bleach and then stratified for 2-3 days at 4ºC. Seeds were then germinated on one-half-strength concentration Murashige and Skoog medium (½ MS) plates containing 0.8% plant agar and 1% sucrose in a growth chamber with 16-hrs light, 8-hrs dark at 22ºC. Tomato *Solanum lycopersicum* cultivar Rio Grande PtoR (*Pto/Pto, Prf/Prf*) and CRISPR *core* lines were sown in agronomy mix in growth chambers under 16/8 hr light/dark cycles, 24°C, about 200 µmol/$m^2$/s light intensity, 50 to 65% humidity, and grown to six to seven-weeks old.

***Measuring ROS Production and Antagonism***

For measuring ROS, leaf punches were taken using cork borer no. 1 (diameter of 4 mm) from equivalent leaves or leaflets from the cotyledon leaves across biological replicates (for csp22, the $5^{th}$ leaflet). Leaf punches (4 per plant) were floated on their abaxial side in 190 µl of sterile water in a 96-well white microtiter plate for 18 to 24 hours in the dark. To test ROS elicitation, sterile water was removed and a 100 µl solution of sterile water, L-012 (Wako Chemicals, Cat. No. 143556-24-5), horseradish peroxidase (HRP) (Sigma Aldridge, Cat. No. P6782), and 100 nM to 1 µM of MAMP peptide dissolved in sterile water or 100% DMSO was added (see SI Appendix Table S2 for solvent used). For elf18 (acyl-MSKEKFERTKPHVNVGTI) and flg22 (QRLSTGSRINSAKDDAAGLQIA) tested on *Arabidopsis,* 20 µM L-012 and 20 ug/mL HPR was used (27). For csp22 tested on tomato, 40 µg/mL HPR was used. For MAMPs which were dissolved in 100% DMSO, an equivalent volume of 100% DMSO was added to the controls. Assay measurements were taken every 1.5 minutes for a minimum 60 minutes. Luminescence was measured using a TriStar LB 941 plate reader (Berthold Technologies). Across all ROS experiments, the maximum RLU was determined for each leaf punch and averaged between punches per plant.

Peptide antagonism: For antagonist assays, the leaf disks were incubated in a solution of water and candidate antagonist MAMP peptide (100 nM to 1 µM) for 18 to 24 hours in the dark. The candidate antagonist MAMP peptide was then removed and the remainder of the assay including elicitation and luminescence measurements are the same as above. Recombinant protein antagonism: Recombinant protein concentrations were quantified and between 100 nM and 500 nM of each protein was added in solution with HRP and L-012. The solution was immediately used for elicitation. Lysate antagonism: Initial protein concentrations were quantified and between 1 and 2.5 µg/mL was added to HPR and L-012 solution and immediately used for elicitation. For all ROS assays, a minimum 12 plants were used. To compare ROS plates, plants and punches per treatment were averaged and scaled based on the appropriate controls.

***Measuring alkalinization and antagonism in Tobacco BY-2 Cells***

The tobacco cell line BY-2 (*Nicotiana tabacum* L. cv. Bright Yellow 2) was culture on a modified MS medium supplemented with 100 µg/mL timentin (32). Callus tissue was passaged every three to four weeks on MS medium with 2% phytoagar. Transplanted calli in petri dishes were wrapped in parafilm and incubated at 25-26°C in the dark. To establish suspension cultures, a 1.5x1.5 cm slice of calli was transferred to 100 mL of modified MS media in a 250 mL flask, covered with autoclavable paper, sealed with a rubber band and wrapped in foil. The flask was incubated for 10 to 14 days in a 26°C shaker rotating at 130 rpm in the dark. Cultures were passed by transferring 10 mL of cells via a stereological pipette into ~90 mL of media. After three passages, multiple cultures were grown in parallel and were grown for experimentation.

To assess alkalinization changes due immune induction, cells were passage from the main line and grown in similar conditions for 5 to 7 days until cell density was about $3.0x10^6$, which was quantified using a hemi-cytometer. Cells were then aliquoted into 20 mL glass flat bottom vials (Thermo Fisher Scientific, Cat. No. 6PCV20-1F) and sealed the tops with Millipore tap (Micropore, Cat. No. 1530-1). The cell vials were then placed on a shaker rotating at 100 rpm at room temperature at ~30° angle such that cells were in continuous motion. After two to three hours of incubation, the pH probe and meter were prepared for measurements similarly to previously established protocols (33). Between 10 nM of peptides were used to measure induced alkalinization and antagonism was measured by first treating with 50 nM candidate antagonist for three minutes, then treating with 10 nM agonist. The pH change was sampled every 10 seconds for up to 30 minutes.

***Plant tissue-based immune assays***

For measuring MAPK induction, *Arabidopsis thaliana* Col-0 and *efr* seeds were germinated on ½ MS plates for five days. The seedlings were then transferred to a 48-well plate (Costar, Cat. No. CLS3548) containing 600 µL of liquid ½ MS medium. Plates were sealed with 3M Micropore Surgical Tape (Micropore, Cat. No. 1530-1) and transferred back to the growth chamber. After nine days of growth, the 14-days-old seedlings were incubated with water or 100 nM of elf18 variants for 0 and 15 min before being pooled for harvest. Three seedlings per treatment were pooled. Seedlings were ground to fine powder in liquid nitrogen and resuspended in 200 µL of extraction buffer 50 mm HEPES (pH 7.5), 50 mm NaCl, 10 mm EDTA, 0.2% Triton X-100, 1× Pierce protease inhibitor mini tablets (Thermo Scientific, Cat. No. A32955), and 1x Pierce phosphatase inhibitor mini tablets (Thermo Scientific, Cat. No. A32957). Total proteins were isolated by centrifugation at 21,000 × g at 4 °C for 10 min and Laemmli buffer was added for subsequent SDS–PAGE. To measure the protein concentration, we used Pierce 660 nm Protein Assay (Thermo Scientific, Cat. No. 22660) with Ionic Detergent Compatibility Reagent (Thermo Scientific, Cat. No. 22663). Equal concentration of protein was loaded per lane and separated by SDS–PAGE. Activated MAPK3 and MAPK6 were detected by immunoblotting using p44/42 MAPK (pERK) antibodies at a concentration of 1:2000 (Cell Signaling Technology, Cat. No. 4370), followed by secondary rabbit antibodies at a concentration of 1:3000 (Biorad, Cat. No. 1705046). Both primary and secondary antibody were suspended in 1x TBS-T with 5% BSA. The experiment was repeated three times with similar results.

For analysis of callose deposition, *Arabidopsis thaliana* Col-0 and *efr* seeds were germinated on ½ MS plates for four days. The seedlings were then transferred to a 48-well plate (Costar, Cat. No. CLS3548) containing 500 µL of liquid ½ MS medium, sealed with 3M Micropore Surgical Tape (Micropore, Cat. No. 1530-1) and transferred back to the growth chamber. Three seedlings were transferred per treatment. After four days of growth, the ½ MS medium was replaced with water or 1 µM of elf18 variants and the seedlings were incubated in a shaker for 18 to 20 h in the growth chamber. Seedlings were fixed in ethanol/acetic acid (3:1, v/v) at 37ºC for about 2 to 3 hours or overnight at room temperature (until cotyledons were completely cleared), with one change of fixing solution. Seedlings were subsequently rehydrated in 70%, 50% and 30% ethanol for 30 min each step. Then the seedlings were washed one time with 150mM $K_2HPO_4$, pH 9.5 and stained with 0.01% (w/v) aniline blue in 150mM $K_2HPO_4$, pH 9.5 for 1 h in dark. The cotyledons were loaded onto the slides with 50% glycerol and callose deposition was imaged by fluorescence microscopy (Leica CMS) using a DAPI filter. Callose quantification was performed with Fiji as previously described (28,29). Experiments were repeated three times with similar results.

To assess seedling growth inhibition (SGI) we followed Gómez-Gómez et al., 1999 with modifications (30). *Arabidopsis thaliana* Col-0 and *efr* seeds were germinated on ½ MS plates for four days. The seedlings were then transferred to a 48-well plate (Costar, Cat. No. CLS3548) containing 500 µL of liquid ½ MS medium supplemented with 100 nM of elf18 variants. Plates were sealed with 3M Micropore Surgical Tape (Micropore, Cat. No. 1530-1) and transferred back to the growth chamber. After eight days of growth, the seedlings were patted dry and weighed. Each experiment included eight seedlings per treatment. A mock control (½ MS medium) and canonical active elf18 sequence (acyl-SKEKFERTKPHVNVGTIG) were assayed on each plate. Experiments were repeated three times with similar results.

Ethylene induction was measured as previously described (31). For measurements of ethylene, each dot represents one measurement (from one tube with water and three leaf pieces); four replicates per treatment. Leaf pieces (squares of approx. 2 x 2-3 mm) were from two individual plants, four- to five-weeks-old, cut the evening before, and incubated on water overnight and induced using 1 µM csp22 variants. Ethylene induction was measured using a Shimadzu Gas Chromatograph GC-14.

***Development of tomato core mutants***

To generate the *core* mutants in the tomato (*Solanum lycopersicum*) cultivar Rio Grande (RG)-PtoR, we designed two guide RNAs (gRNA1: 5'- GTAGCATTTGACAATGTCCC-3'; gRNA2: 5'-GACTGGCCTGGGGTCTCATG-3') that target the first exon of *CORE* using the software Geneious R11. The gRNA cassette was cloned into the p201N:Cas9 binary vector as described previously (34,35). Tomato transformation was performed at the Biotechnology Center at the Boyce Thompson Institute as described previously (36). Mutations were confirmed by Sanger sequencing at the Biotechnology Resource Center (BRC) at Cornell University.

***Bacterial Strains and Growth conditions***

*Clavibacter michiganensis* strains were grown in tryptone broth with yeast (TBY) medium at 28°C, *Escherichia coli* was grown in Luria broth (LB) medium at 37°C, *Streptomyces spp.* ND05-3B was grown on yeast malt extract (YME) at 28°C, and *Pseudomonas syringae* pv. *tomato* strains were grown in (NYGB) medium at 28°C. The following concentrations were used for antibiotic selection: 50 µg/mL gentamycin, 25 µg/mL kanamycin, 25 µg/mL chloramphenicol, 50 µg/mL spectinomycin, and 100 µg/mL rifamycin.

**Assessing syntenic gene structure in *Clavibacter***

Seven genomes across five species in the genome were assessed for their gene structure surrounding the CSP loci identified in the MAMP mining pipeline. Those selected included *Clavibacter michiganensis* NCPPB382 (Accession: GCF_000063485.1), *C. michiganensis* Z001 (Accession: GCF_002931335.1), *C. capsici* PF008 (Accession: GCF_001280205.1), *C. insidiosus* ATCC10253 (Accession: GCF_003076355.1), *C. nebraskensis* NCPPB2581 (Accession: GCF_000355695.1), and *C. tessellarius* ATCC33566 and CFBP8017, respectively (Accession: GCF_002240635.1 and GCF_002151185).1 Briefly, the region within each genome was identified via a blast search and regions of interest were extracted via a custom Biopython script. The extracted regions in Genbank format were fed into clinker, a gene clustering program (v0.0.27) (37). The outputs clustered gene structures were output as an SVG file and loci color was manually edited in Inkscape (v1.0.2).

***Measuring expression of CSPs in Clavibacter michiganensis***

To assess if all CSP loci are expressed, we cultured the type-strain Gram-positive pathogen *C. michiganensis* NCPPB382 *in vitro* in both rich TBY medium and minimal m9 medium supplemented with xylem sap, which mimics the xylem vasculature (38). To collect xylem sap, tomatoes were grown for 6 weeks in chamber conditions described above and a similar procedure was followed (39). Briefly, using scissors sterilized with 70% ethanol, the stem was cut about 10-12 cm above the cotyledons and additional leaflets were removed. Wounds were sealed with parafilm. After re-sterilizing the scissors, a P1000 tip was cut such that the diameter of the tip can go on top the stem. The tips were parafilmed to the stem and a small bag placed over the tip. Over the next two hours, xylem sap was collected. The collected sap was filter sterilized after passing through a 0.22 µM filter.

*C. michiganensis* NCPPB382 was grown on TBY agar plates. After three days, the plates were scraped, and the bacteria transferred to 25 mL flasks of TBY broth for a starter culture. The starter cultures were grown at 28°C for ~20 hours and used to inoculate growth curves. Two flasks of m9 broth with xylem sap were inoculated with *C. michiganensis* NCPPB382 to a starting $OD_{600}$ of 1.0 and two flasks of TBY broth were inoculated with *C. michiganensis* NCPPB382 to a starting $OD_{600}$ of 0.5. One mL of culture was removed from each flask at 6 hours and 24 hours of growth. The flasks were grown shaking at 28°C.

One mL samples were removed from cultures pelleted via centrifugation. 500 µL of supernatant was then removed and two volumes of RNA Protect Bacteria Reagent (QIAGEN, California, USA) was added. The cultures were resuspended, incubated at room temperature for five minutes, and pelleted again. They were then frozen with liquid nitrogen and transferred to a -80°C freezer until RNA extraction. RNA was extracted with the SV Total RNA isolation kit (Promega, Wisconsin, USA). Growth curves and RNA extraction were repeated three times. All PCR primers are listed in Table S3.

RNA was quantified by Qubit via a high sensitivity RNA kit (Thermo Fisher Scientific) and the concentration was standardized for each repeat. Random primers were annealed to the RNA and first-strand synthesis performed with M-MLV reverse transcriptase (Promega, Wisconsin, USA). For qPCR, cDNA was diluted 1:5 and SsoFast EvaGreen Supermix was used (BioRad, California, USA). *BipA* was used as a housekeeping gene (40). Each repeat was performed in a single qPCR plate. Except for rich media TBY broth at 6 hours, where only one sample was used due to space constraints, two samples were used for each broth and time point. qPCR measurements were detected on a Bio-Rad CFX96 Real-Time PCR Detection System.

***Collection of bacterial lysates***

An initial starter 20 mL culture of *C. michiganensis* NCPPB382 was incubated at 28°C at 200 rpm until the optical density ($OD_{600}$) equals one. A subculture was made in 300 mL TBY medium and cultured in similar conditions until the optical density ($OD_{600}$) equaled two. *Pseudomonas syringae* pv. *tomato* DC3000Δ*flaA* was cultured similarly in NYGB. For *Streptomyces spp.* ND05-3B, spores were populated on YME medium over four to five days at 28°C (41). Using a cell scraper and sterile water, spores were collected in 25 mL and quantified to a concentration of 9.05 x$10^6$ spores/mL via a hemacytometer. Cells were spun down at 5000 rpm and resuspended in lysis buffer (300 mM NaCl, 50 mM Hepes, pH 7.0) with 100 mM phenylmethylsulfonyl fluoride (PMSF) (Thermo Scientific) and 0.2 U/mL DnaseI (Thermo Scientific). The chilled cells were sheered via an Emusiveflex-C3 High-Pressure Homogenizer (Avestin, Ottawa, Ontario, Canada) until the solution turned clear. The collected lysates were passed through a 0.22-micron filter, froze with liquid nitrogen and store in the -80°C until used. To test induction of immunity by ROS production, protein lysates were quantified using a Pierces 660 assay using BSA as a standard control and tested using the same procedure as describe in Plant tissue-based immune assays at 1 μg/mL concentration.

***Expression and Purification of Clavibacter CSPs***

The coding sequence of each CSP from NCPPB382 was codon-optimized for *E. coli* and synthesized into entry vector, pENTRY, from either ThermoFisher Scientific or Twist Biosciences (Waltham, Massachusetts or San Francisco, California, USA). Each vector was transformed into *E. coli* DH5a and after subsequent colony isolation and plasmid miniprep, each vector sequence was confirmed via whole plasmid sequencing (Plasmidsaurus, Eugene, Oregon, USA). Each CSP sequence was then transferred into expression vector pRSET via LR clonase using manufacturer's instructions (Thermo Scientific). The subsequent reactions were transformed into *E. coli* DH5a and putative transformants were checked via PCR for the insert and the sequence was confirmed via whole plasmid sequencing. Subsequent CSP expression vectors were transformed into *E. coli* expression genotype BL21(DE3).

For protein expression, confirmed vector-carrying transformants were grown in 50 mL LB media at 37°C until $OD_{600}$ at least 1. The initial culture was used to seed a larger culture of LB medium, which was grown until $OD_{600}$ 0.4 and 0.6, at which a final concentration of 0.5 mM IPTG was added to each culture and shaken at 200 rpms at 28°C for four hours. Bacterial cells were pelleted by centrifugation at 4°C and resuspended in buffer A (20 mM Tris, 10 mM Imidazole, 500 mM NaCl, pH 8.0) with 1 mM PMSF, 10 uM leupeptin serine protease inhibitor, and 10 ug/mL lysozyme. Cell suspension was incubated on ice for 30 minutes, then sonicated in four 10 second cycles at 30% output and 30% duty cycle with 15 second breaks between each cycle. Cells were centrifuged at 5,000 rpm for 20 minutes at 4°C. The supernatant was incubated with prepped Ni-NTA Resin (Thermo Fisher Scientific Cat. No. 88222), previously washed 3x with buffer A, for one hour at 4°C on a rotating shaker. The protein-bound beads were centrifuged at 5,000 rpm for 10 minutes at 4°C, washed 3x with buffer A, then incubated with elution buffer B (20 mM Tris, 500 mM NaCl, 5% Glycerol, pH 8.0). Each incubating elution set had increasing concentrations of Imidazole from 50 to 250 mM. The eluted protein was collected after the beads were centrifuged at 5,000 rpm for 10 minutes at 4°C. At each step from cell fraction to elution, sample was collected and prepped to run on a 15% SDS-Page gel. The gel was either stained with Coomassie SimpleBlue SafeStain (Thermo Fisher Scientific, Cat. No. LC6060) or transferred to a PVDF membrane and blotted with 1:2000 anti-6xHis-HRP antibody in 5% BSA-TBST (Thermo Fisher Scientific, Cat. No. MA1-21315-HRP). Purified recombinant protein was desalted via 3.5K MWCO dialysis cassettes (Thermo Fisher Scientific, Cat. No. A52966) in buffer (20 mM Tris, 50 mM NaCl, 5% glycerol, pH 8.0) overnight at 4°C and concentration quantified via Qubit protein assay kit (Thermo Fisher Scientific, Cat. No. Q33211).

***Overexpression of cspB and transformation into Pst via Tri-parental mating***

A *Pst* DC3000 codon-optimized gBlock of *Cm* NCPPB382 *cspB* was synthesized (Azenta, USA) and amplified via PCR with primers carrying restriction sites NdeI and EcoRI (Thermo Fisher Scientific). Both board host range plasmid pDSK519-ngfp and amplified fragment were digested with *Nde*I and *Eco*RI and ligated using T4 ligase (Thermo Fisher Scientific) (42). Standard DH5a *E. coli* transformation was performed and plasmid was confirmed via whole plasmid sequencing (Plasmidsaurus). For pDSK519-*cspB*, an additional restriction with *Bsi*WI and ligation was performed to move the chloramphenicol antibiotic cassette from a Gateway cassette into pDSK519.

Cultures of donor vectors, helper plasmids (pRK2013/pRK600), and recipient strains were grown overnight in no antibiotics (43,44). Cultures were mixed in the following ratio (1 Donor: 1 Helper: 5 Recipient) and spot plated onto a NYGB plate with no antibiotics and incubated in 28°C. After two days, spots were re-streaked onto NYGB plates which contained antibiotics for selection of recipient strain with donor plasmid and grown overnight. Transformants were confirmed for the donor plasmid via standard gDNA preparation and PCR screening.

Overexpression of *cspB* was confirmed via in vitro growth in NYGB media, protein lysates were separated using standard 15% SDS-Page Gel, probed using an anti-HA-HRP conjugated antibody (Cat. No. 26183-HRP, Thermo Fisher Scientific). In detail, cultures were grown to an OD of 1.0, cells were spun down and resuspended in water and 3x Laemmli buffer supplemented with DDT. Cells incubated at 95°C for five minutes, briefly put on ice and spun at max speed. Proteins samples were run on an 15% SDS-PAGE Gel, transferred to a nitrocellulose membrane via semi-dry, and blotted with 1:1000 anti-HA-HRP conjugated antibody in blocking solution (Thermo Fisher Scientific). Chemiluminescence was detected using the Bio-Rad Chemidoc system. Protein loading for *in vitro* bacterial expression was visualized using the SilverQuest Silver Straining Kit (Thermo Fisher Scientific).

**Disease assays and bacterial titers of *P. syringae* in tomato**

Tomato Rio Grande (RG)-PtoR and mutant line *core* were grown to about seven-weeks of age just before flowering. Three days before inoculation, strains *Pto* DC3000ΔΔΔ carrying empty vector pDSK519-ngfp or constitutive expressing vector pDSK519-CO-*cspB*-HA (pDSK519-cspB) were streaked from glycerol stocks and cultured on NYGB plates with appropriate antibiotics. Cells were scraped from the plate, resuspended in plain NYGB and an inoculum was adjusted to an $OD_{600}$ = 0.00001 (or $1x10^4$ CFUs/mL) by resuspension in 5 mM $MgCl_2$. Several leaves per plant were infiltrated with a needless syringe and a black sharpie was used to circle the infiltration area. After three days post inoculation, using a #3 cork borer, a single punch was taken from each plant with four plants per genotype per treatment and moved to a 1.5 mL tube containing 200 µl 5 mM $MgCl_2$. The tissues were ground and dilutions were carried out between $10^{-2}$ and $10^{-5}$ and plated on NYGB plates containing ½ antibiotics and 50 µg/mL cycloheximide. After 2 days incubation at 28°C, colonies were counted and CFU per mg tissue was determined.

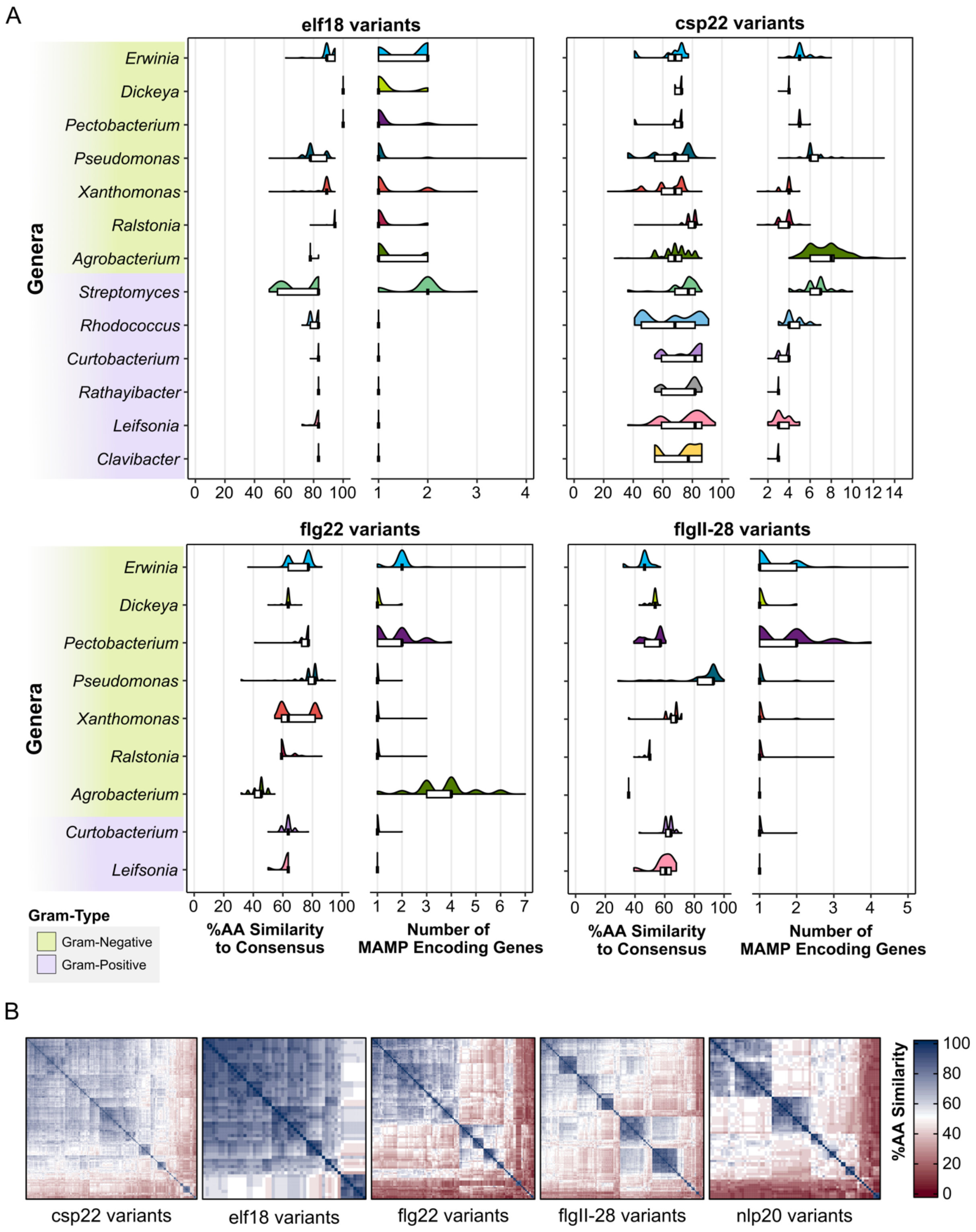

**Fig. S1. Copy number and conservation differ across on immunogenic features and bacterial genera. (A)** Violin plots of percent amino acid (AA) similarity of epitope variants in comparison to each respective consensus sequence across all genera sampled. Tukey's boxplots are plotted on top. **(B)** All-by-all amino acid (AA) similarity comparisons for each MAMP independent of the consensus sequence.

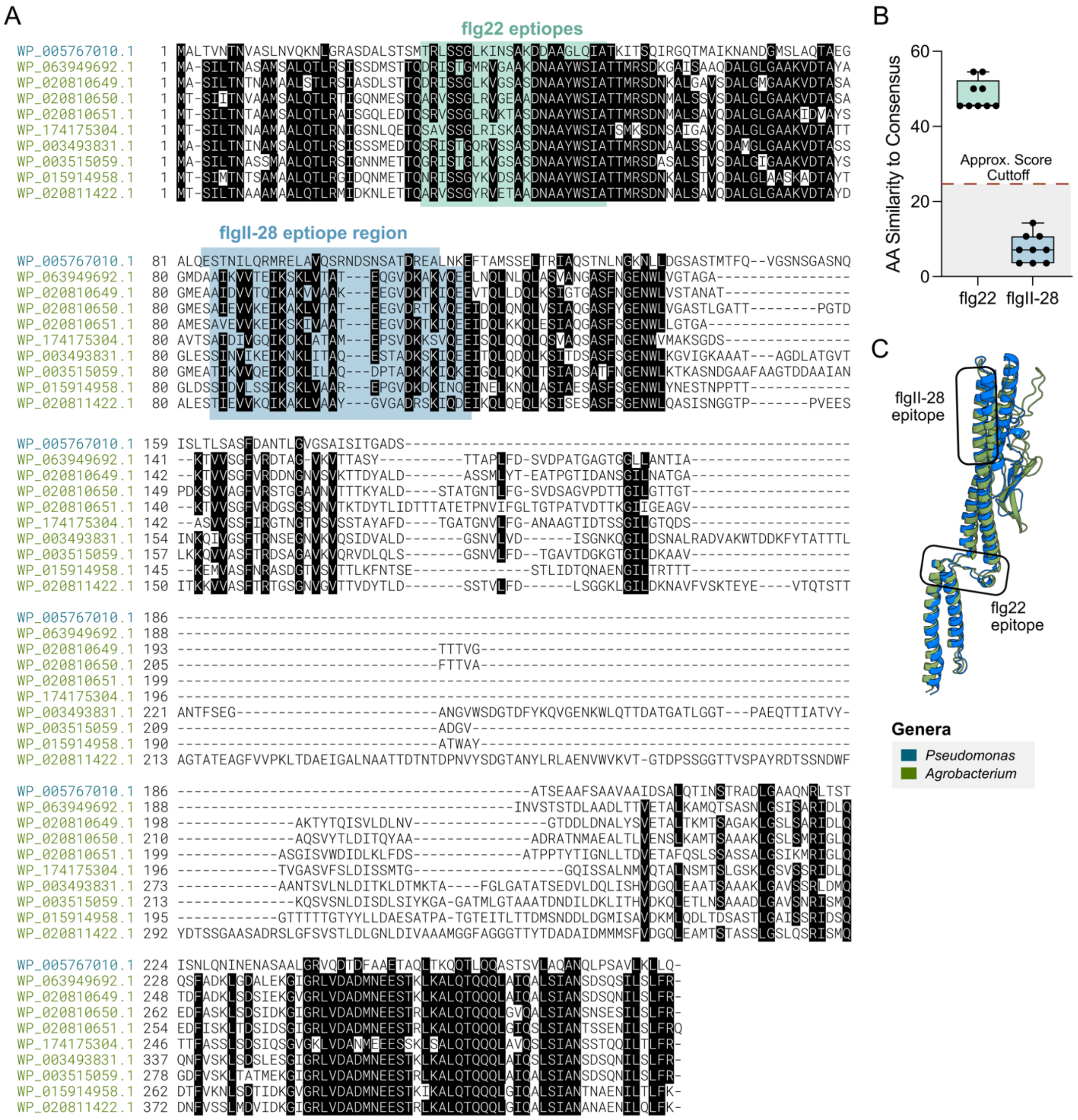

**Fig. S2. Examples of divergent FliC proteins with undetectable flgII-28 epitopes. (A)** Alignment of flagellin proteins from the reference *Pseudomonas syringae* pv. *tomato* DC3000 and *Agrobacterium,* which carry a flg22 epitope but fail detection for flgII-28. **(B)** Percent amino acid (AA) similarity of epitope variants from (a) in comparison to each respective consensus sequence. **(C)** Structural modeling of flagellin from *Pto* DC3000 (blue = WP_005767010.1) and *Agrobacterium* (green = WP_020810649.1). RMSD = 1.579.

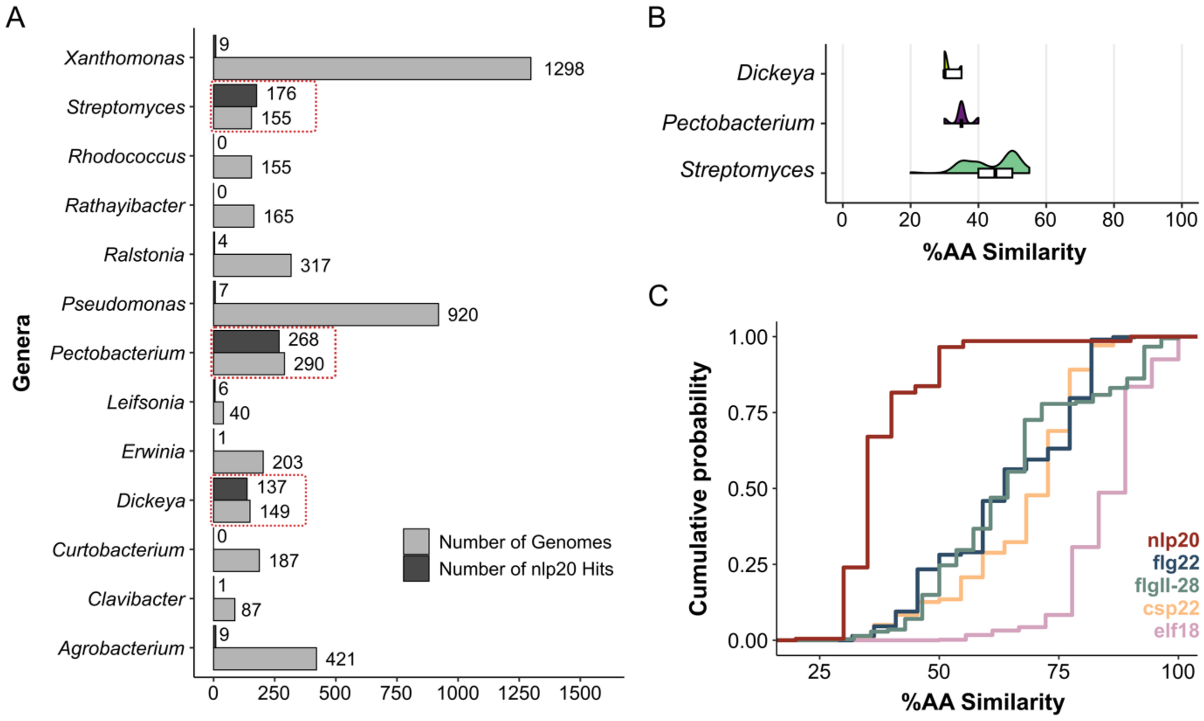

**Fig. S3. Nlp20 is present but not conserved across diverse bacteria. (A)** The number of nlp20 hits in respect to the number of genomes analyzed. Red boxes highlight genera where nlp20 MAMPs are abundantly conserved. **(B)** Violin plot with Tukey's boxplot on top of percent AA similarity of nlp20 epitopes in comparison to the consensus sequence across the three major genera's nlp20 are found. **(C)** Cumulative probability of Fig. 1C in respect to percent amino acid (AA) similarity.

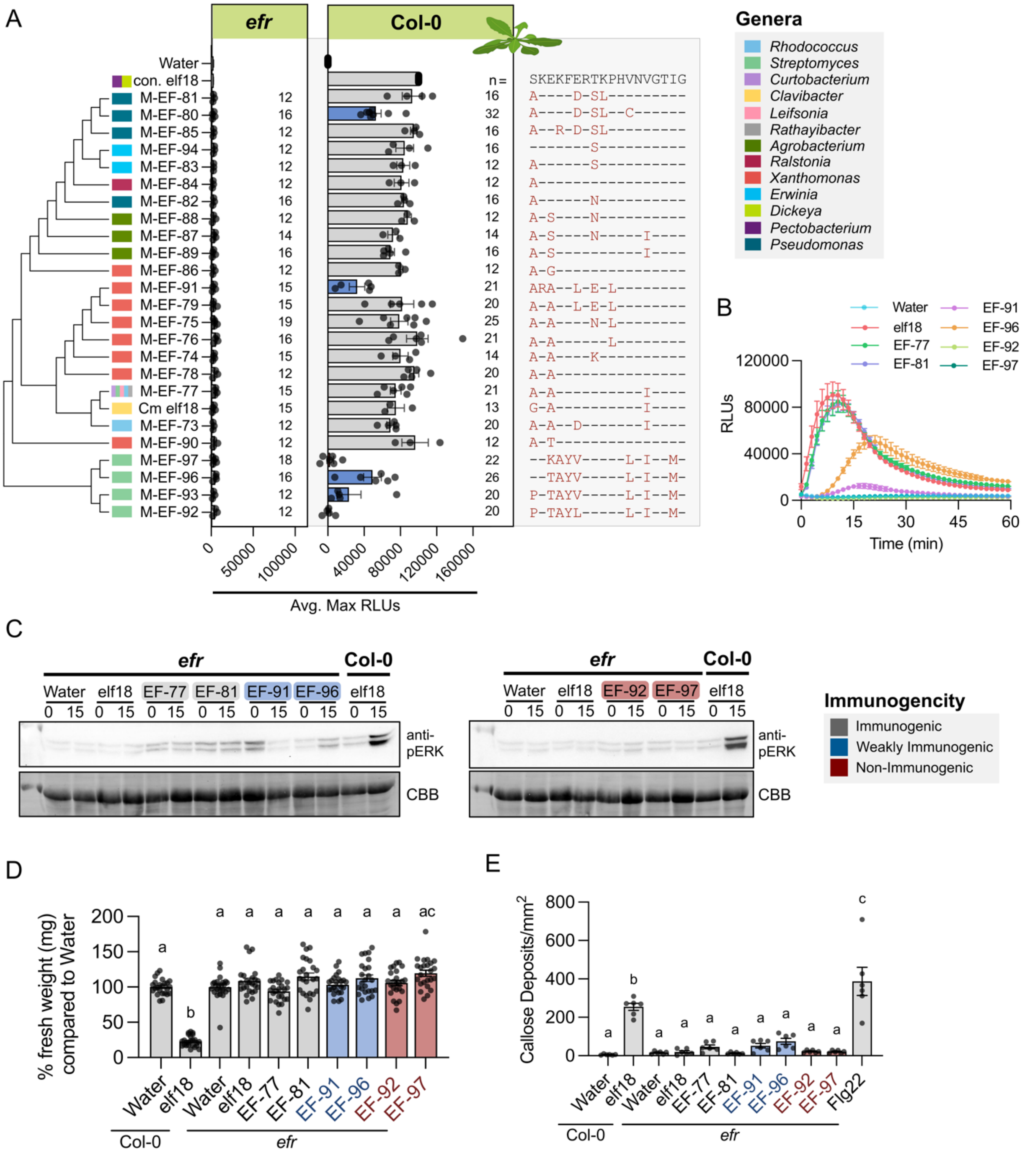

**Fig. S4. Perception and specificity of elf18 variants. (A)** Left: Cladogram based on an alignment of elf18 epitope sequences with tips labeled by genera. Middle: ROS induction of elf18 variants (100 nM concentration) in *Arabidopsis* Col-0 and the *efr* mutant line. Each data point represents an average max reactive light unit (RLU) from 4 plants with 4 leaf disks per plant. The number of plants sampled are plotted to the right. For induction in Col-0, values of each plate were adjusted to a scale of 0 to 100000 maximum RLUs based on the controls (water and consensus elf18; also referred to as con. elf18 and elf18). A one-way ANOVA and Tukey's mean comparison was used to determine significance on non-scaled data, p<0.05

(grey = not significantly different from the consensus; blue = significantly less than the consensus but higher than water; red = not significantly different from water). Across all plates, water and consensus elf18 were tested alongside *efr* transgenic plants. Error bars = SEM. Right: Alignment of epitopes tested with changes from the consensus noted in red. **(B)** Independent analysis of ROS induction of a subset of elf18 variants. Error bars = SEM. Two plants per plate across six plates was conducted. All peptides were tested at 100 nM. **(C)** Induction of MAPK by elf18 variants (100 nM) in *Arabidopsis* Col-0 and the *efr* mutant at zero- and 15-minutes post infiltration. CBB = protein loading. **(D)** *Arabidopsis thaliana* seedling growth inhibition by either water (mock) or elf18 variants (100 nM). One point = 1 plant. Eight plants per biological replicate. The experiment was complete three times. A one-way ANOVA and Tukey's mean comparison was used to determine significance. **(E)** Quantified deposition of callose in *Arabidopsis* Col-0 and *efr* mutant by water and elf18 variants (1 μM). Values are from one representative experiment which includes an average of at least two images per leaf from two leaves per plant, three plants per treatment. The experiment was repeated three times with similar results. A one-way ANOVA and Tukey's mean comparison was used to determine significance for panel (D) and (E), $p<0.0001$.

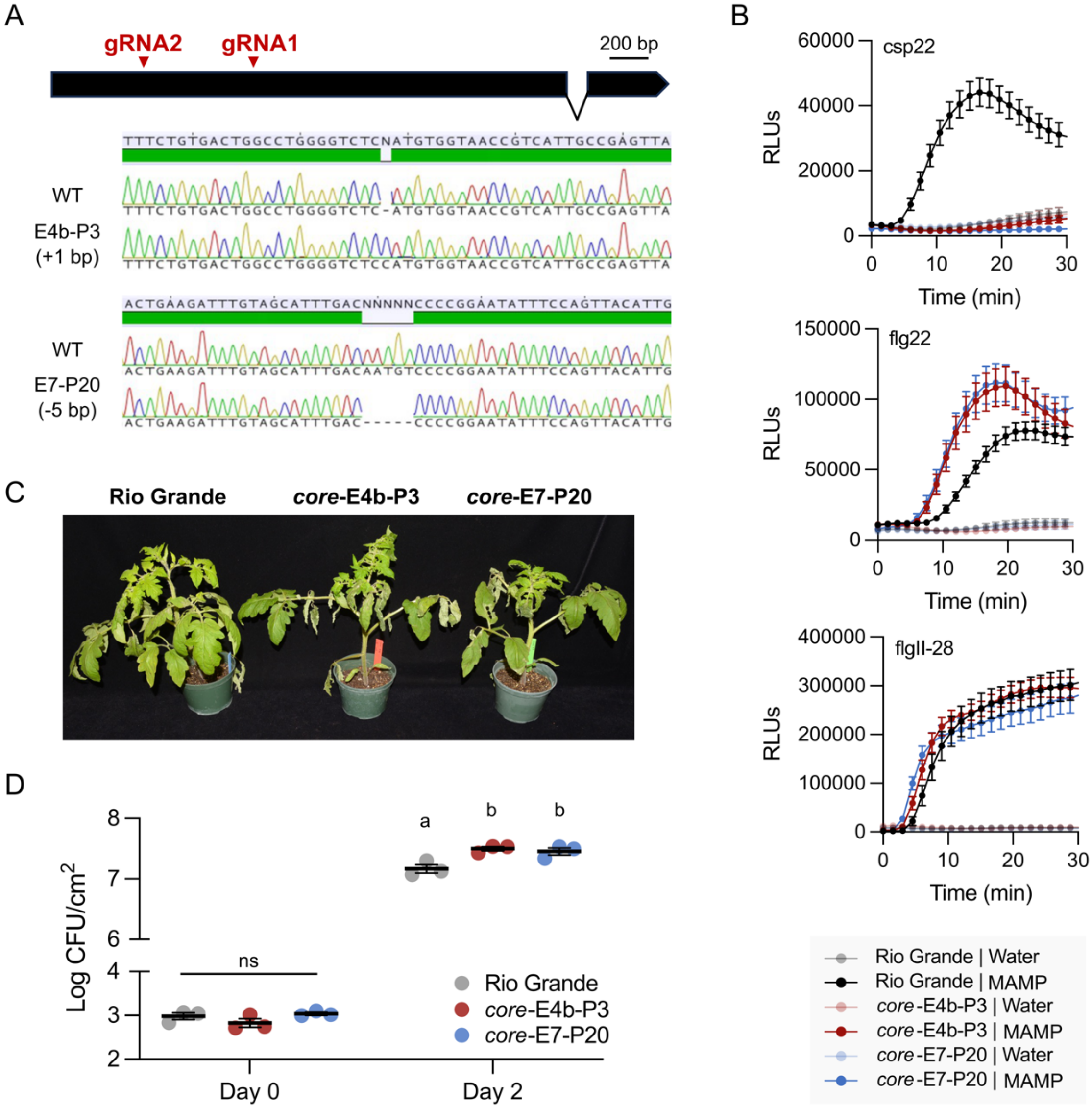

**Fig. S5. Development of *core* mutant lines via CRISPR-cas9. (A)** Gene structure of *CORE* (Solyc03g096190) with the region targeted for edits denoted in red. Bottom: Sanger sequencing reads of targeted edits in two independent lines compared to wild-type (WT) Rio Grande (RG)-PtoR. **(B)** ROS production in response to consensus MAMP peptides csp22 (200 nM), flg22 (100 nM), and flgII-28 (100 nM) in two mutant lines. **(C)** The *core* mutants were more susceptible to *Pseudomonas syringae* pv. *tomato* DC3000Δ*avrPto*Δ*avrPtoB*Δ*flaA* (DC3000ΔΔΔ). Four-week-old core mutants and wild-type RG-PtoR plants were vacuum infiltrated with 5 x $10^4$ cfu/mL DC3000ΔΔΔ*.* Photographs of disease symptoms were taken five DPI. **(D)** Bacterial populations in leaves were measured at three hours (Day 0) and two days (Day 2) after infiltration. Error bars = SEM. Different letters indicate significant differences based on a one-way ANOVA followed by Student's t test ($p < 0.05$). ns = no significant difference. Three plants for each genotype were tested per experiment. The experiment was performed twice with similar results.

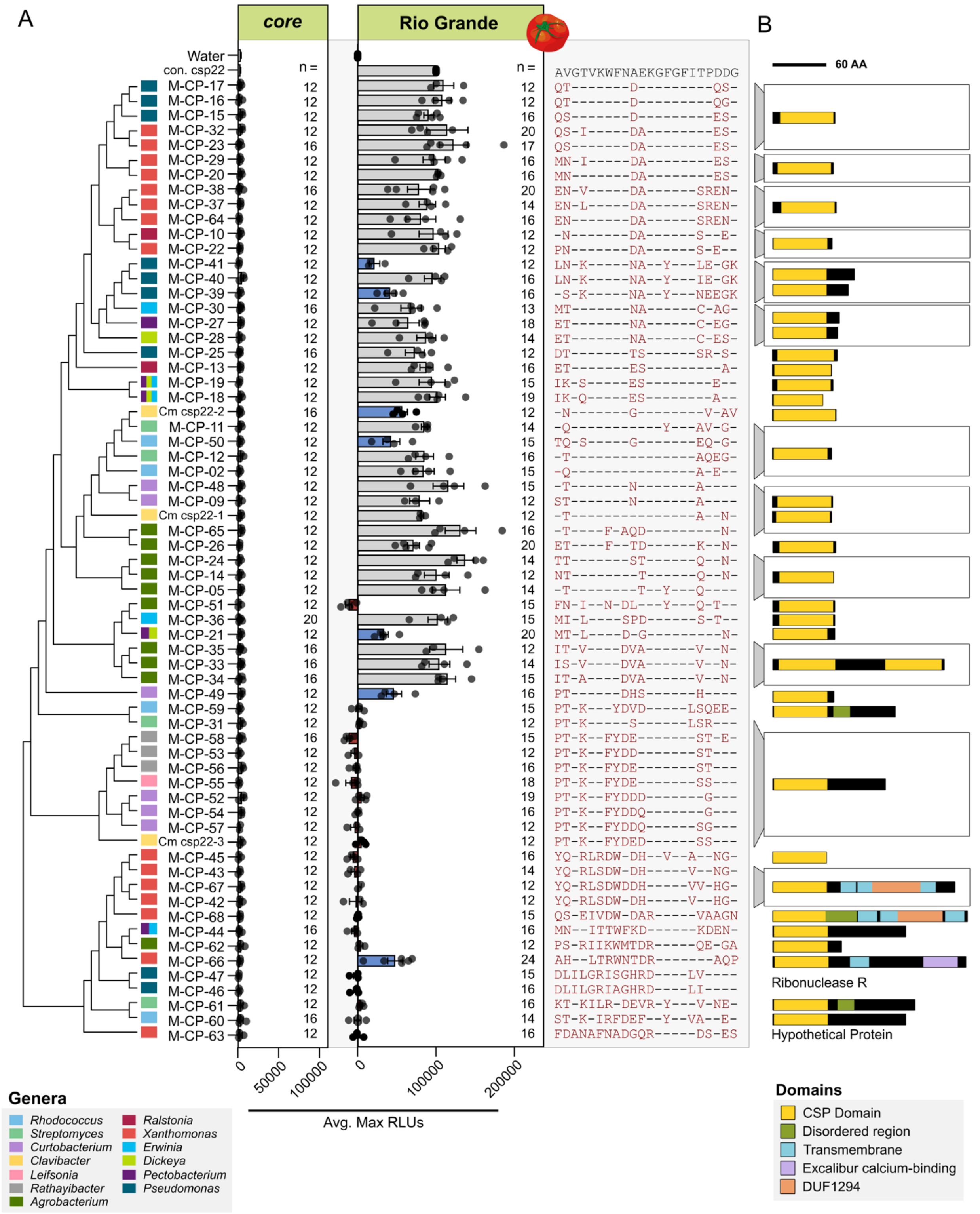

**Fig. S6: csp22 variants exhibit differential immune induction in tomato. (A)** Left: Cladogram based on alignment of epitope sequences with tips label by genera. Middle: ROS induction of csp22 variants (200 nM concentration) in tomato Rio Grande and a *core* mutant line. Each data point represents an average max RLU from 4 plants with 4 leaf disks per plant. The number of plants sampled are plotted to the right. For immune induction in Rio Grande, values of each plate were adjusted to a scale of 0 to 100000 maximum RLUs based on the controls (water and consensus csp22/con. csp22). A one-way ANOVA and Tukey's mean comparison was used to determine significance on non-scaled data, $p<0.05$ (grey = not significantly different from the consensus; blue = significantly less than the consensus but higher than water; red = not significantly different from water). Across all plates, water and con. csp22 were tested. Error bars = SEM. Right: Alignment of epitopes tested with changes from the consensus noted in red. **(B)** Domain structure of diverse CSPs derived from csp22 epitopes in (A) predicted using InterProScan. For CSPs with two domains, the epitope from the first domain was tested.

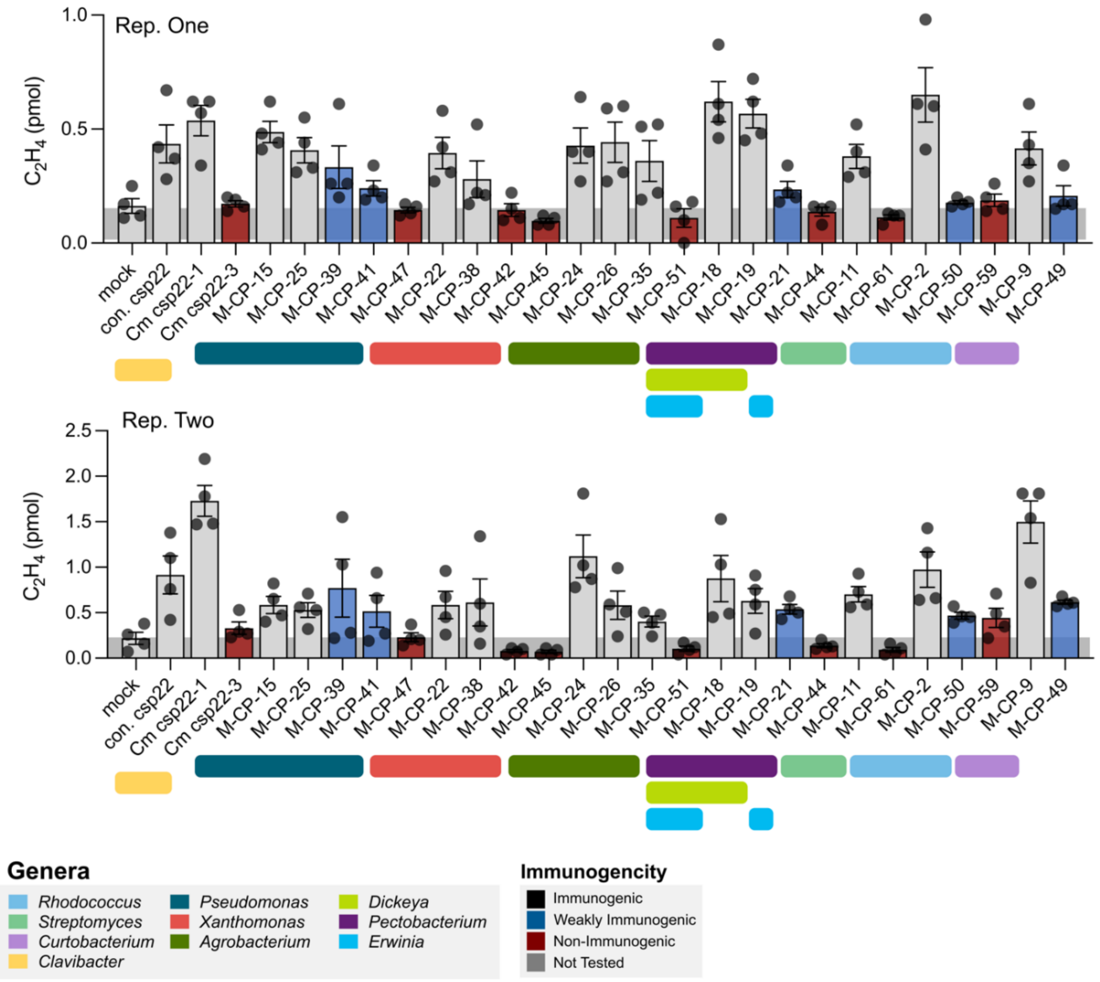

**Fig. S7. Ethylene production induced by csp22 variants.** Induction of ethylene by csp22 variants (1 µM) in 4- to 5-week-old tomatoes. Each dot represents one measurement (from one tube with water and 3 leaf pieces); four replicates per treatment. Bars are colored to match immunogenicity conclusions from ROS data in Figure 4A (grey = not significantly different from the consensus; blue = significantly less than the consensus but higher than water; red = not significantly different from water). Error bars = SEM.

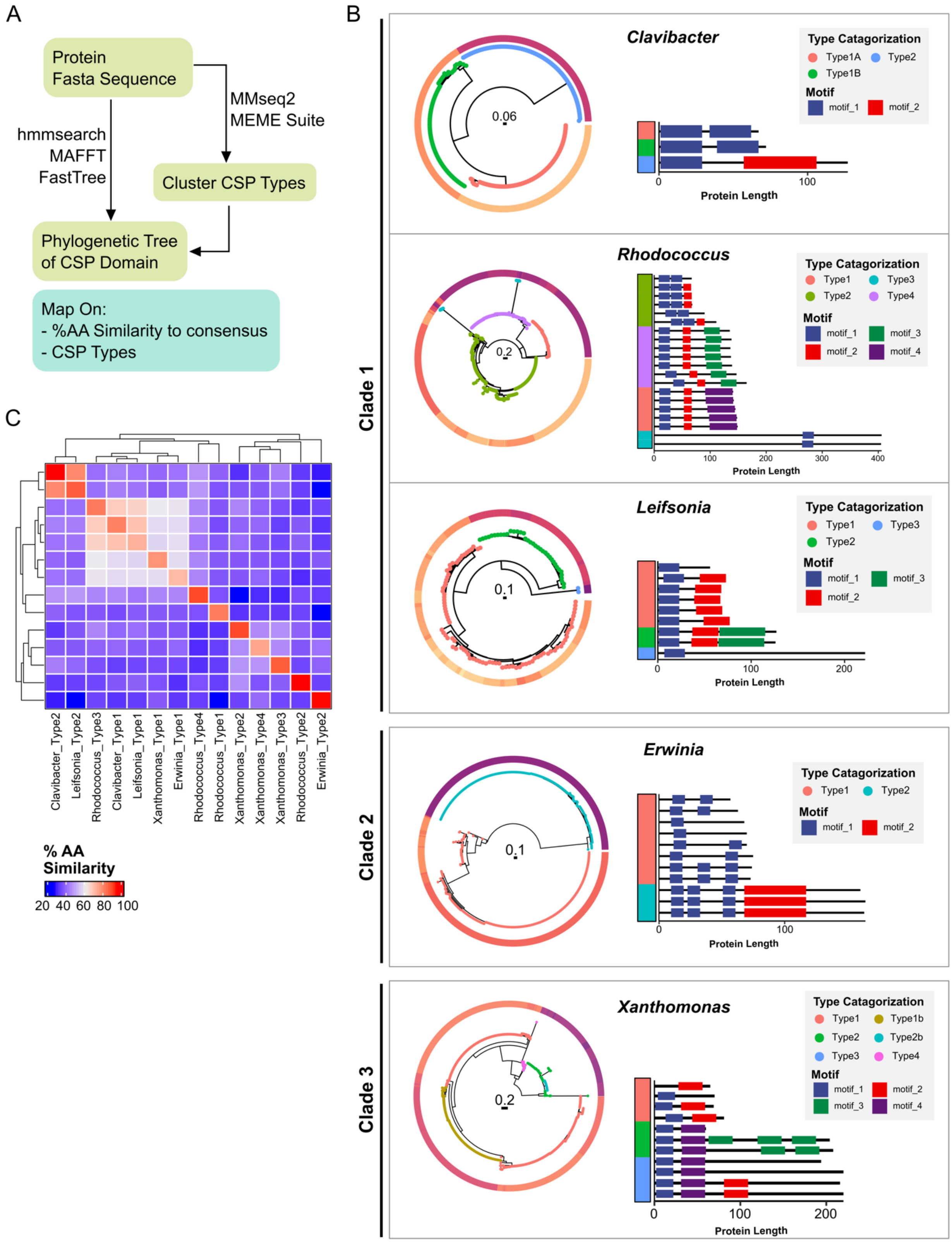

**Fig. S8. Classification of CSP Types. (A)** Pipeline for CSP classification. **(B)** Phylogenetic trees of cold shock proteins in different genera. Across all trees, tips are labeled by type categorization which is based on clade structure, mmseq2 cluster classification, and motif structure predicted by MEME. The motif classification of represented cold shock proteins can be seen on the right of each tree and similarly labeled by the same type classification. Genera assessed include those found in clades highlighted in Figure 3. **(C)** BlastP assessment of CSP typing across genera. Those CSP types whose average similarity are above 70% are predicted as orthologs.

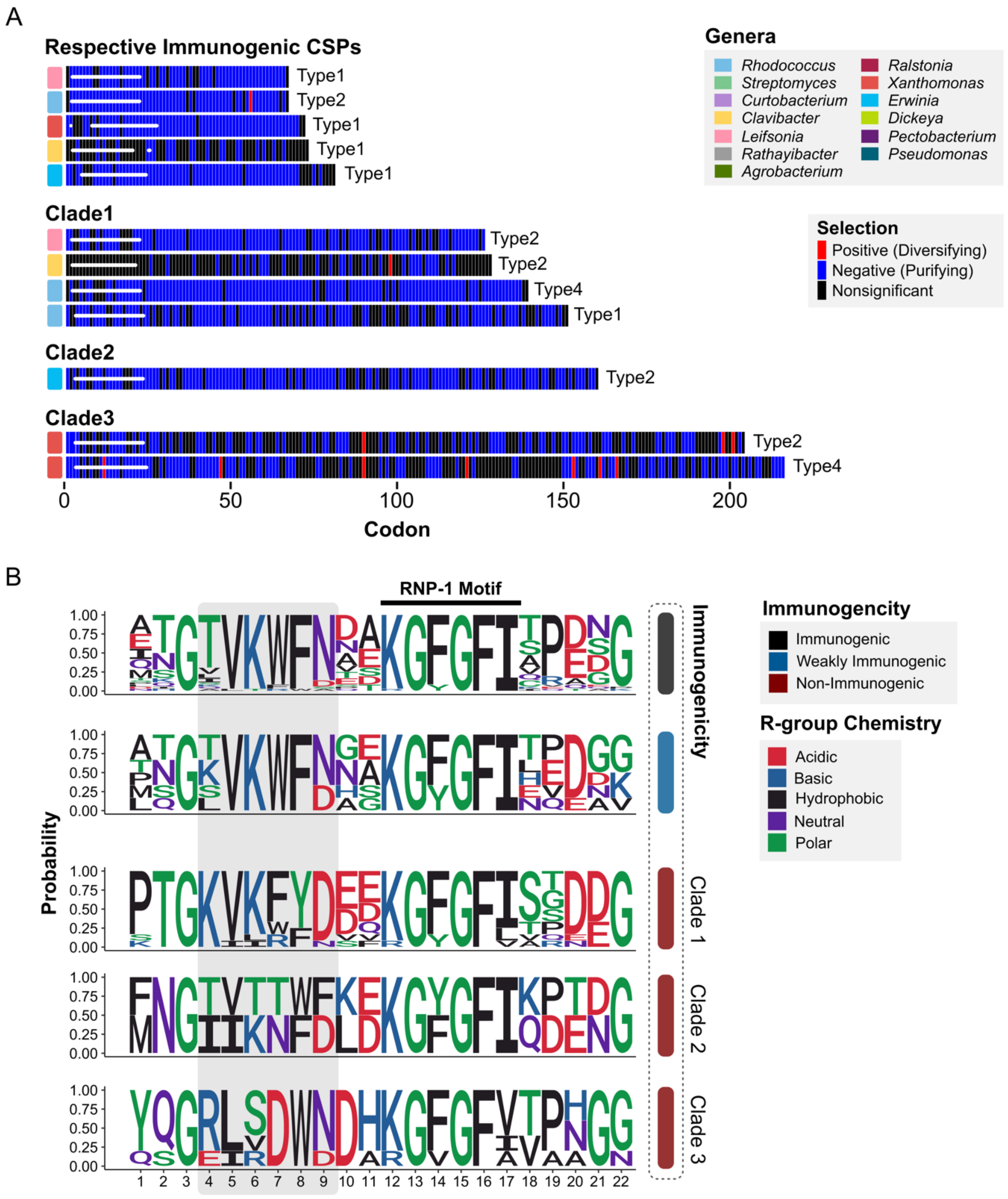

**Fig. S9. Convergently evolved non-immunogenic CSPs are under purifying selection. (A)** $d_N/d_S$ tests of immunogenic (top) and non-immunogenic CSP types (Clades 1 - 3) across multiple bacterial genera. Sites which are significant by Bayesian approximation for selection are colored in blue (negative, purifying) or red (positive, diversifying). The region of the epitope is represented in the white bar. Typing of CSPs is described in Supplemental Figure 8. **(B)** Weblogos of immunogenic, weakly immunogenic, and non-immunogenic csp22 variants from each major clade tested by ROS and shown in Figure 3.

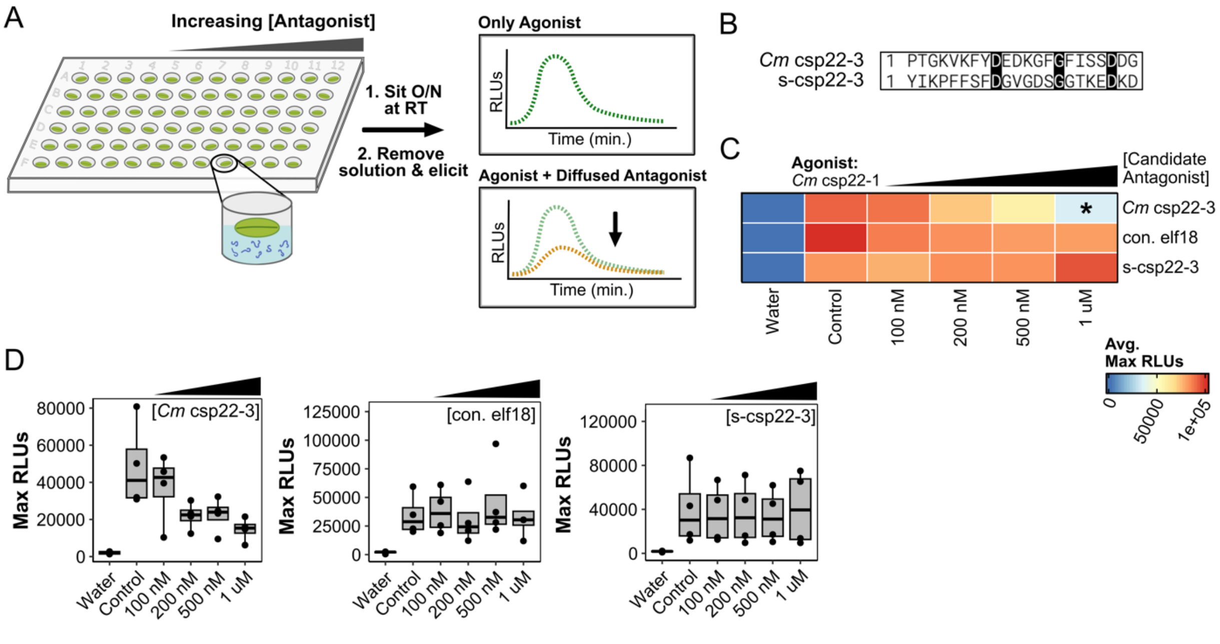

**Fig. S10. Non-immunogenic *Cm* csp22-3 antagonizes CORE perception. (A)** Diagram of the assay to assess MAMP antagonism. Concentrations for antagonism were 100 nM, 200 nM, 500 nM, and 1 μM (left to right). Immunogenic agonist was tested at 200 nM. All antagonism assays included four punches per plant, four plants for each experiment, and the experiment was repeated at least three times. **(B)** Alignment of *Cm* csp22-3 with a scrambled version of the peptide (s-csp22-3). **(C)** *Cm* csp22-3 assessed for antagonism against immunogenic *Cm* csp22-1. ROS screen for antagonism in Rio Grande tomato. Control denotes positive control (untreated agonist) and concentrations listed are of the candidate antagonist. Maximum RLU averages were adjusted to a scale of 0 to 100,000 based on the controls (water and untreated agonist). The assay was also repeated using con. elf18 and s-csp22-3. A one-way ANOVA and Tukey's mean comparison was used to determine significance in respect to the untreated agonist control (denoted *: $p < 0.05$). **(D)** Representative assays for (C).

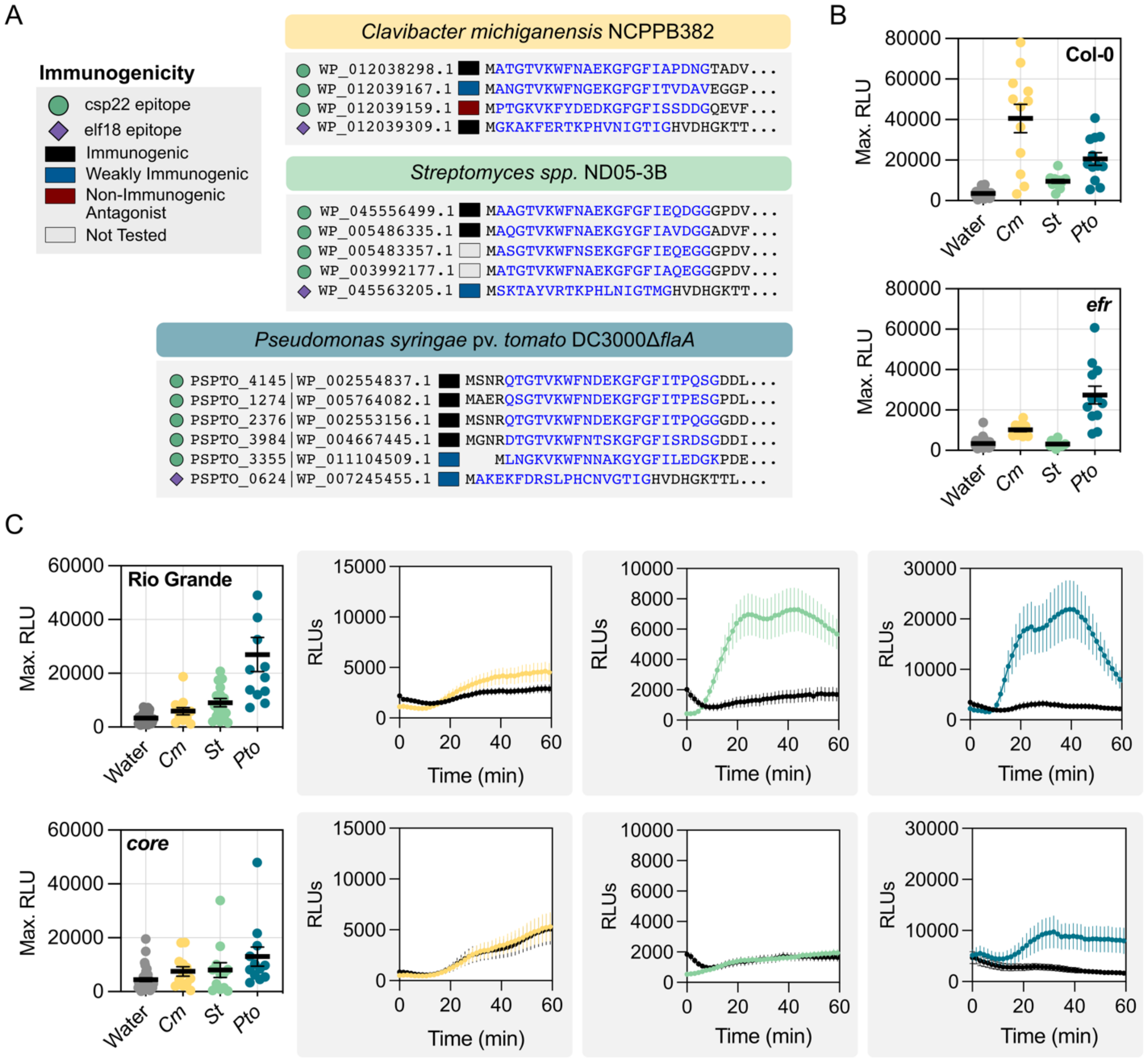

**Fig. S11. MAMP perception from bacterial lysates on *Arabidopsis* and tomato. (A)** Predicted csp22 and elf18 epitopes (highlighted in blue) encoded in bacterial genomes and their immunogenic outcomes based on the ROS screen. **(B and C)** ROS induction of bacterial lysates (1 µg/mL) in (B) *Arabidopsis* Col-0 and the *efr* mutant and (C) tomato Rio Grande and the *core* mutant. *Cm* = *C. michiganensis* NCPPB382, *St* = *Streptomyces spp.* ND05-3B, and *Pto* = *Pseudomonas syringae* pv. *tomato* DC3000Δ*flaA*. Consensus flg22 (100 nM) was used as a positive control (not shown). Each dot represents an average of the maximum RLUs of four disks per plant (n = 12). Representative plates shown on the right. Error bars = SEM.

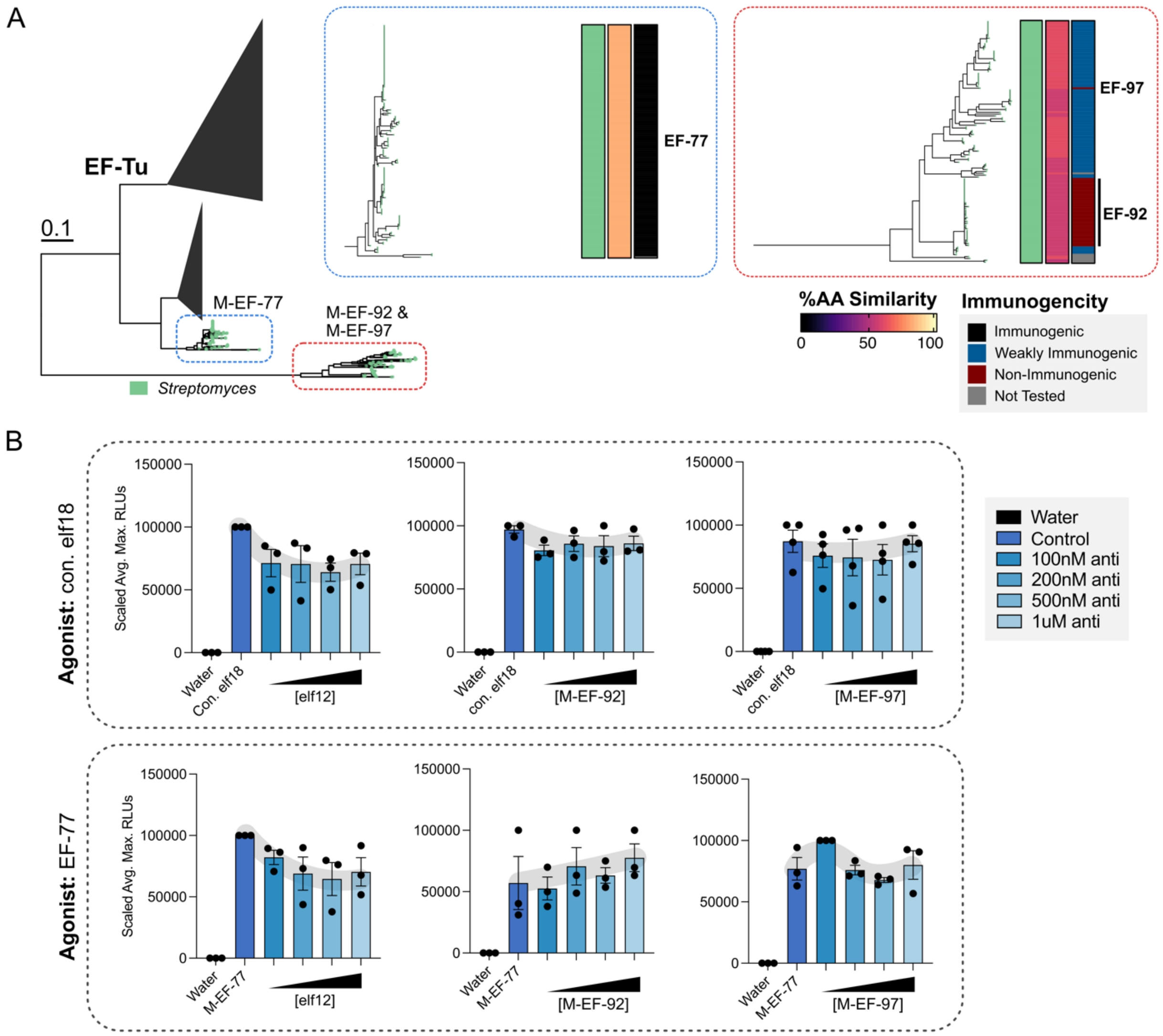

**Fig. S12. No antagonism was identified for elf18. (A)** Subset of the EF-Tu protein tree from Figure 2 highlighting those from *Streptomyces.* The immunogenic variant (M-EF-77) is found in the clade outlined in blue and the non-immunogenic variants (M-EF-92 and M-EF-97) are found in the clade outlined in red. **(B)** ROS screen for antagonism in *A. thaliana* Col-0. Control denotes positive control (untreated agonist) and concentrations listed are of the candidate antagonist tested. Maximum RLU averages were adjusted to a scale of 0 to 100000. Error bars = SEM. Elf12 has been previously reported as a weak antagonist.

**Table S1: MAMPs detected from the computational pipeline in this study.** Hits detected via BlastP and Local Alignment were combined. Duplicates were then removed before final filtering for partial proteins and off-targets based on the corresponding conserved motif.

| MAMP | MAMPs Detected via BlastP | MAMPs Detected via Local Alignment | Total MAMPs Detected Post Filtering |
|---|---|---|---|
| Elf18 | 4660 | 3960 | 4404 |
| Csp22 | 20665 | 19851 | 20542 |
| Flg22 | 3217 | 4958 | 5042 |
| FlgII-28 | 3697 | 3419 | 3661 |
| Nlp20 | 124 | 636 | 613 |

**Table S2. Custom Synthetic Peptides used in this study.** Epitope sequence colored: green = immunogenic, blue = weakly immunogenic (predicted/confirmed deviant), orange = non-immunogenic (bolded, antagonist), not colored = not tested. Solvents colored: pink = water, yellow = 100% DMSO. Frequency = number of occurrences across all mined epitopes.

| Reference Name | Sequence | Frequency | Modification | Purity | Solvent | Company | Notes |
|---|---|---|---|---|---|---|---|
| M-CP-1 | ANGTVKWFNAEKGFGFITVDGG | 175 | None | | | Biomatik, Cambridge, Ontario, Canada | Unable to be synthesized |
| M-CP-2 | AQGTVKWFNAEKGFGFIAPEDG | 151 | | 95.22 | Water | | |
| M-CP-3 | ANGTVKWFNAEKGYGFITVDGG | 129 | | | | | Unable to be synthesized |
| M-CP-4 | ATGTVKWFNAEKGFGFIEQDGG | 145 | | | | | Unable to be synthesized |
| M-CP-5 | ATGTVKWFNATKGYGFIQPDDG | 240 | | 96.35 | Water | | |
| M-CP-6 | ATGTVKWFNDAKGFGFITPDEG | 251 | | | | | Unable to be synthesized |
| M-CP-7 | ATGTVKWFNETKGFGFITPDGG | 261 | | | | | Unable to be synthesized |
| M-CP-8 | PTGTVKWFNSEKGFGFIAPDDG | 142 | | | | | Unable to be synthesized |
| M-CP-9 | STGTVKWFNNEKGFGFIAPDDG | 109 | | 95.11 | Water | | |
| M-CP-10 | ANGTVKWFNDAKGFGFISPDEG | 153 | | 96.7 | Water | | |
| M-CP-11 | AQGTVKWFNAEKGYGFIAVDGG | 153 | | 95.78 | Water | | |
| M-CP-12 | ATGTVKWFNAEKGFGFIAQEGG | 127 | | 96.17 | Water | | |
| M-CP-13 | ETGTVKWFNESKGFGFITPDAG | 140 | | 95.8 | Water | | |
| M-CP-14 | NTGTVKWFNATKGFGFIQPDNG | 219 | | 96.5 | Water | | |
| M-CP-15 | QSGTVKWFNDEKGFGFITPESG | 860 | | 95.33 | Water | | |
| M-CP-16 | QTGTVKWFNDEKGFGFITPQGG | 852 | | 95.69 | Water | | |
| M-CP-17 | QTGTVKWFNDEKGFGFITPQSG | 857 | | 95.99 | Water | | |
| M-CP-18 | IKGQVKWFNESKGFGFITPADG | 578 | | 95.88 | Water | | |
| M-CP-19 | IKGSVKWFNESKGFGFITPEDG | 596 | | 95.51 | Water | | |
| M-CP-20 | MNGTVKWFNDAKGFGFITPESG | 127 | | 95.35 | Water | | |
| M-CP-21 | MTGLVKWFDAGKGFGFITPDNG | 275 | | 96.26 | Water | | |
| M-CP-22 | PNGTVKWFNDAKGFGFISPEDG | 1148 | | 95.32 | Water | | |
| M-CP-23 | QSGTVKWFNDAKGFGFITPESG | 483 | | 95.54 | Water | | |
| M-CP-24 | TTGTVKWFNSTKGFGFIQPDNG | 337 | | 95.15 | Water | | |
| M-CP-25 | DTGTVKWFNTSKGFGFISRDSG | 876 | | 96.21 | Water | | |
| M-CP-26 | ETGTVKFFNTDKGFGFIKPDNG | 245 | | 96.1 | Water | | |
| M-CP-27 | ETGTVKWFNNAKGFGFICPEGG | 275 | | 96.53 | Water | | |
| M-CP-28 | ETGTVKWFNNAKGFGFICPESG | 126 | | 96.97 | Water | | |
| M-CP-29 | MNGIVKWFNDAKGFGFITPESG | 205 | | 97.06 | Water | | |
| M-CP-30 | MTGTVKWFNNAKGFGFICPAGG | 132 | | 96 | Water | | |
| M-CP-31 | PTGKVKWFNSEKGFGFLSRDDG | 144 | | 95.56 | Water | | |
| M-CP-32 | QSGIVKWFNDAKGFGFITPESG | 101 | | 95.01 | Water | | |
| M-CP-33 | ISGVVKWFDVAKGFGFIVPDNG | 130 | | 95.38 | DMSO | | |
| M-CP-34 | ITGAVKWFDVAKGFGFIVPDNG | 124 | | 95.56 | DMSO | | |
| M-CP-35 | ITGVVKWFDVAKGFGFIVPDNG | 444 | | 95.37 | DMSO | | |
| M-CP-36 | MIGLVKWFSPDKGFGFISPTDG | 131 | | 95.91 | DMSO | | |
| M-CP-37 | ENGLVKWFNDAKGFGFISRENG | 249 | | 96.53 | Water | | |
| M-CP-38 | ENGVVKWFNDAKGFGFISRENG | 824 | | 95.22 | Water | | |
| M-CP-39 | ASGKVKWFNNAKGYGFINEEGK | 115 | | 95.39 | Water | | |
| M-CP-40 | LNGKVKWFNNAKGYGFIIEDGK | 118 | | 95.93 | Water | | |

| | | | | | | |
|---|---|---|---|---|---|---|
| M-CP-41 | LNGKVKWFNNAKGYGFILEDGK | 270 | | 95.56 | Water | |
| M-CP-42 | YQGRLSDWNDHKGFGFVTPHGG | 326 | | 96.04 | Water | |
| M-CP-43 | **YQGRLSDWNDHKGFGFVTPNGG** | 219 | | 97.17 | Water | |
| M-CP-44 | MNGTITTWFKDKGFGFIKDENG | 438 | | 95.03 | Water | |
| M-CP-45 | YQGRLRDWNDHKGVGFATPNGG | 198 | | 95.11 | Water | |
| M-CP-46 | DLILGRIAGHRDGFGFLIPDDG | 109 | | 95.85 | DMSO | |
| M-CP-47 | DLILGRISGHRDGFGFLVPDDG | 710 | | 95.63 | DMSO | |
| M-CP-48 | ATGTVKWFNNEKGFGFIAPDDG | 58 | | 95.53 | Water | |
| M-CP-49 | PTGTVKWFDHSKGFGFIHPDDG | 59 | | 96.68 | Water | |
| M-CP-50 | TQGSVKWFNGEKGFGFIEQDGG | 91 | | 95.41 | Water | |
| M-CP-51 | **FNGIVKNFDLEKGYGFIQPTDG** | 62 | | 95.47 | Water | |
| M-CP-52 | PTGKVKFYDDDKGFGFITGDDG | 100 | | 96.1 | Water | |
| M-CP-53 | **PTGKVKFYDDEKGFGFISTDDG** | 31 | | 95.26 | Water | |
| M-CP-54 | PTGKVKFYDDQKGFGFITGDDG | 50 | | 95.97 | Water | |
| M-CP-55 | PTGKVKFYDEEKGFGFISSDDG | 11 | | 96.26 | Water | |
| M-CP-56 | PTGKVKFYDEEKGFGFISTDDG | 85 | | 95.94 | Water | |
| M-CP-57 | PTGKVKFYDDQKGFGFISGDDG | 19 | | 95.65 | Water | |
| M-CP-58 | PTGKVKFYDEEKGFGFISTDEG | 31 | None | 95.48 | Water | |
| M-CP-59 | **PTGKVKWYDVDKGFGFLSQEEG** | 88 | | 97.33 | Water | |
| M-CP-60 | STGKVIRFDEFKGYGFVAPDEG | 84 | | 97.19 | Water | |
| M-CP-61 | KTGKILRFDEVRGYGFIVPNEG | 50 | | 97.11 | Water | |
| M-CP-62 | PSGRIIKWMTDRGFGFIQEDGA | 6 | | 95.34 | Water | |
| M-CP-63 | FDANAFNADGQRGFGFIDSDES | 5 | | 95.71 | DMSO | |
| M-CP-64 | ENGTVKWFNDAKGFGFISRENG | 82 | | 97.11 | Water | |
| M-CP-65 | ATGTVKFFAQDKGFGFITPDNG | 95 | | 95.87 | Water | |
| M-CP-66 | AHGTLTRWNTDRGFGFITPAQP | 15 | | 95.5 | Water | |
| M-CP-67 | YQGRLSDWDDHKGFGFVVPHGG | 56 | | 96.49 | Water | |
| M-CP-68 | QSGEIVDWNDARGFGFIVAAGN | 1 | | 95.57 | DMSO | |
| M-EF-73 | AKAKFDRTKPHVNIGTIG | 72 | | 97.31 | Water | Biomatik, Cambridge, Ontario, Canada |
| M-EF-74 | AKAKFERKKPHVNVGTIG | 37 | | 96.87 | Water | |
| M-EF-75 | AKAKFERNKLHVNVGTIG | 24 | | 96.19 | Water | |
| M-EF-76 | AKAKFERTKLHVNVGTIG | 19 | | 95.77 | Water | |
| M-EF-77 | AKAKFERTKPHVNIGTIG | 582 | | 97.08 | Water | |
| M-EF-78 | AKAKFERTKPHVNVGTIG | 998 | | 97.45 | Water | |
| M-EF-79 | AKAKFLREKLHVNVGTIG | 54 | | 96.95 | Water | |
| M-EF-80 | AKEKFDRSLPHCNVGTIG | 102 | | 96.39 | Water | |
| M-EF-81 | AKEKFDRSLPHVNVGTIG | 522 | | 97.17 | Water | |
| M-EF-82 | AKEKFERNKPHVNVGTIG | 253 | | 97.19 | Water | |
| M-EF-83 | AKEKFERSKPHVNVGTIG | 124 | | 96.6 | Water | |
| M-EF-84 | AKEKFERTKPHVNVGTIG | 312 | | 97.36 | Water | |
| M-EF-85 | AKERFDRSLPHVNVGTIG | 13 | N-terminal acetylation | 96.57 | Water | |
| M-EF-86 | AKGKFERTKPHVNVGTIG | 83 | | 97.35 | Water | |
| M-EF-87 | AKSKFERNKPHVNIGTIG | 360 | | 97.18 | Water | |
| M-EF-88 | AKSKFERNKPHVNVGTIG | 62 | | 95.03 | Water | |
| M-EF-89 | AKSKFERTKPHVNIGTIG | 18 | | 98.6 | Water | |
| M-EF-90 | AKTKFERTKPHVNVGTIG | 70 | | 95.65 | Water | |
| M-EF-91 | ARAKFLREKLHVNVGTIG | 42 | | 95.74 | Water | |
| M-EF-92 | PKTAYLRTKPHLNIGTMG | 38 | | 95.76 | Water | |
| M-EF-93 | PKTAYVRTKPHLNIGTMG | 28 | | 95.83 | Water | |
| M-EF-94 | SKEKFERSKPHVNVGTIG | 78 | | 95.72 | Water | |
| M-EF-95 | SKEKFERTKPHVNVGTIG | 330 | | 96.41 | Water | |
| M-EF-96 | SKTAYVRTKPHLNIGTMG | 60 | | 95.6 | Water | |
| M-EF-97 | SKKAYVRTKPHLNIGTMG | 1 | | 95.85 | Water | |
| Cm elf18 | GKAKFERTKPHVNIGTIG | 63 | | 96.9 | Water | Genscript, Piscataway, New Jersey, USA |
| Cm csp22-1 | ATGTVKWFNAEKGFGFIAPDNG | 87 | None | 99.2 | Water | |
| Cm csp22-2 | ANGTVKWFNGEKGFGFITVDAV | 72 | | 96.2 | DMSO | |

| | | | | | |
|---|---|---|---|---|---|
| Cm csp22-3 | **PTGKVKFYDEDKGFGFISSDDG** | 87 | | 95.5 | Water |
| s-csp22-3 | YIKPFFSFDGVGDSGGTKEDKD | N/A | | 95.3 | Water |
| elf12 | **SKEKFERTKPHV** | N/A | N-terminal acetylation | 96.7 | Water |

**Table S3: Oligonucleotides used in this study.**

| Name | Product Size (bp) | Sequence | Tm (°C) | Reference |
|---|---|---|---|---|
| Primers for amplification of *Clavibacter CSP*s and screening of tomato *core* mutants. | | | | |
| Name | Product Size (bp) | Sequence | Tm (°C) | Reference |
| Amplify_cspB_CO_Pst_F | 408 | GGTGGTcatatgATGCCCACGG | 60 | This Study |
| Amplify_cspB_CO_Pst_R | | GGTGGTgaattcTCAAGCATAGTC | | |
| Screen_cspB_pDSK519_F | 441 | GGTGGTcatatgATGCCCACGG | 60 | |
| Screen_cspB_pDSK519_R | | GGTGGTgaattcTCAAGCATAGTC | | |
| Amplify_cmx_from_pselact_ko_w_promotor_BsiWI_RS_F | 827 | GGTGGTCGTACGtatccagtcactatggcggcc | 56 | |
| Amplify_cmx_from_pselact_ko_BsiWI_RS_R | | GGTGGTCGTACGTTAGTAAGCCGGATCCTCTAGA | | |
| Screen_antibiotic_cassette_pDSK519_F | 1995/1064 | GGGCTATGTGCAACGGGAAT | 60 | |
| Screen_antibiotic_cassette_pDSK519_R | | GCGATCTGGCTATCGCGG | | |
| Primers for screening of codon-optimized *Clavibacter* CSPs in *E. coli* for recombinant protein expression | | | | |
| Name | Product Size (bp) | Sequence | Tm (°C) | Reference |
| attB1_F | Varies | GGGGACAAGTTTGTACAAAAAAGCAGGCT | 57 | This Study |
| attB1_R | | GGGGACCACTTTGTACAAGAAAGCTGGGT | | |
| Primers to screen for bp edits within regions of CORE targeted by gRNA | | | | |
| Name | Product Size (bp) | Sequence | Tm (°C) | Reference |
| Screen_CORE_CRISPR_edit_F | 1184 | ATGATTCTCCCAAAGAATTCTCACTT | 55 | This Study |
| Screen_CORE_CRISPR_edit_R | | ACTTGATTATAAGACAACGAAAGCTC | | |
| Primers for qPCR primer for relative expression in *Cm* NCPPB382 | | | | |
| Name | Product Size (bp) | Sequence | Tm (°C) | Reference |
| NCPPB382_cspA1_ver1_qPCR_F | 125 | TTAGAGCGGGCGGATGTTC | 59 | This Study |
| NCPPB382_cspA1_ver1_qPCR_R | | TGTTCGCTCACTACTCCGC | | |
| NCPPB382_cspA2_qPCR_F | 80 | GCCGAGTAGTGGACGAAGAC | 59 | |
| NCPPB382_cspA2_qPCR_R | | GAGAAGGGGTTCGGGTTCAT | | |
| NCPPB382_cspB_qPCR_F | 51 | CCGAGCTGATGAACCCGAA | 59 | |
| NCPPB382_cspB_qPCR_R | | GCAAGGTGAAGTTCTACGACG | | |
| NCPPB382_bipA_qPCR_F | 63 | GGGTGCTGGTCGTCGTA | 59 | Jiang et al., 2019 |
| NCPPB382_bipA_qPCR_R | | CGAGCCGCTGTTCAAG | | |

#### SI References

1. C. Camacho, *et al.,* BLAST+: architecture and applications. *BMC Bioinform.* 10, 421 (2009).
2. C. Jain, L. M. Rodriguez-R, A. M. Phillippy, K. T. Konstantinidis, S. Aluru, High throughput ANI analysis of 90K prokaryotic genomes reveals clear species boundaries. *Nat Commun* 9, 5114 (2018).
3. D. Stevens, A. Moreno-Pérez, A. Weisberg, C. Ramsing, J. Fliegmann, N. Zhang, M. Madrigal, G. Martin, A. Steinbrenner, G. Felix, G. Coaker, Data from "**Evolutionary dynamics of proteinaceous MAMPs reveals intrabacterial antagonism of plant immune perception.**" Zenodo. Available at http://dx.doi.org/10.5281/zenodo.10724865 (2024).
4. Z. Gu, R. Eils, M. Schlesner, Complex heatmaps reveal patterns and correlations in multidimensional genomic data. *Bioinformatics* 32, 2847–2849 (2016).
5. Z. Gu, L. Gu, R. Eils, M. Schlesner, B. Brors, circlize implements and enhances circular visualization in R. *Bioinformatics* 30, 2811–2812 (2014).
6. O. Wagih, ggseqlogo: a versatile R package for drawing sequence logos. *Bioinformatics* 33, 3645–3647 (2017).
7. S. Xu, *et al.,* Use ggbreak to Effectively Utilize Plotting Space to Deal With Large Datasets and Outliers. *Front. Genet.* 12, 774846 (2021).
8. M. D. Lee, GToTree: a user-friendly workflow for phylogenomics. *Bioinformatics* 35, btz188 (2019).
9. K. Katoh, D. M. Standley, MAFFT Multiple Sequence Alignment Software Version 7: Improvements in Performance and Usability. *Mol Biol Evol* 30, 772–780 (2013).
10. Johnson, L.S., Eddy, S.R., and Portugaly, E. (2010). Hidden Markov model speed heuristic and iterative HMM search procedure. BMC Bioinform. *11*, 431. 10.1186/1471-2105-11-431.
11. B. Q. Minh, *et al.,* IQ-TREE 2: New Models and Efficient Methods for Phylogenetic Inference in the Genomic Era. *Mol Biol Evol* (2020) doi:10.1093/molbev/msaa015.
12. M. Steinegger, J. Söding, MMseqs2 enables sensitive protein sequence searching for the analysis of massive data sets. *Nat Biotechnol* 35, 1026–1028 (2017).
13. T. L. Bailey, J. Johnson, C. E. Grant, W. S. Noble, The MEME Suite. *Nucleic Acids Res.* 43, W39–W49 (2015).
14. X. Li, L. Ma, X. Mei, Y. Liu, H. Huang, ggmotif: An R Package for the extraction and visualization of motifs from MEME software. *Plos One* 17, e0276979 (2022).
15. S. L. Nystrom, D. J. McKay, Memes: A motif analysis environment in R using tools from the MEME Suite. *PLoS Comput. Biol.* 17, e1008991 (2021).
16. M. N. Price, P. S. Dehal, A. P. Arkin, FastTree 2 – Approximately Maximum-Likelihood Trees for Large Alignments. *Plos One* 5, e9490 (2010).
17. K. P. Schliep, phangorn: phylogenetic analysis in R. *Bioinformatics* 27, 592–593 (2011).

18. L.-G. Wang, *et al.,* treeio: an R package for phylogenetic tree input and output with richly annotated and associated data. *Mol. Biol. Evol.* 37, 599–603 (2019).

19. G. Yu, D. K. Smith, H. Zhu, Y. Guan, T. T. Lam, ggtree: an r package for visualization and annotation of phylogenetic trees with their covariates and other associated data. *Methods Ecol. Evol.* 8, 28–36 (2017).

20. S. Xu, *et al.,* ggtreeExtra: Compact Visualization of Richly Annotated Phylogenetic Data. *Mol. Biol. Evol.* 38, 4039–4042 (2021).

21. H.-G. Drost, A. Gabel, I. Grosse, M. Quint, Evidence for Active Maintenance of Phylotranscriptomic Hourglass Patterns in Animal and Plant Embryogenesis. *Mol. Biol. Evol.* 32, 1221–1231 (2015).

22. V. Ranwez, E. J. P. Douzery, C. Cambon, N. Chantret, F. Delsuc, MACSE v2: Toolkit for the Alignment of Coding Sequences Accounting for Frameshifts and Stop Codons. *Mol. Biol. Evol.* 35, 2582–2584 (2018).

23. L.-T. Nguyen, H. A. Schmidt, A. von Haeseler, B. Q. Minh, IQ-TREE: A Fast and Effective Stochastic Algorithm for Estimating Maximum-Likelihood Phylogenies. *Mol. Biol. Evol.* 32, 268–274 (2015).

24. B. Murrell, *et al.,* FUBAR: A Fast, Unconstrained Bayesian AppRoximation for Inferring Selection. *Mol. Biol. Evol.* 30, 1196–1205 (2013).

25. A. Larsson, AliView: a fast and lightweight alignment viewer and editor for large datasets. *Bioinformatics* 30, 3276–3278 (2014).

26. C. Zipfel, *et al.,* Perception of the Bacterial PAMP EF-Tu by the Receptor EFR Restricts Agrobacterium-Mediated Transformation. *Cell* 125, 749–760 (2006).

27. S. Lacombe, *et al.,* Interfamily transfer of a plant pattern-recognition receptor confers broad-spectrum bacterial resistance. *Nat Biotechnol* 28, 365 (2010).

28. J. Schindelin, *et al.,* Fiji: an open-source platform for biological-image analysis. *Nat. Methods* 9, 676–682 (2012).

29. K. N. Mason, G. Ekanayake, A. Heese, Chapter 10 Staining and automated image quantification of callose in Arabidopsis cotyledons and leaves. *Methods Cell Biol.* 160, 181–199 (2020).

30. L. Gómez-Gómez, G. Felix, T. Boller, A single locus determines sensitivity to bacterial flagellin in Arabidopsis thaliana. *Plant J.* 18, 277–284 (1999).

31. M. Albert, *et al.* Arabidopsis thaliana Pattern Recognition Receptors for Bacterial Elongation Factor Tu and Flagellin Can Be Combined to Form Functional Chimeric Receptors*. *J. Biol. Chem.* 285, 19035–19042 (2010).

32. F. Kumagai-Sano, T. Hayashi, T. Sano, S. Hasezawa, Cell cycle synchronization of tobacco BY-2 cells. *Nat. Protoc.* 1, 2621–2627 (2006).

33. G. Fiorin, A. Sánchez-Vallet, B. Thomma, G. Pereira, P. Teixeira, MAMP-triggered Medium Alkalinization of Plant Cell Cultures. *BIO-Protoc.* 10, e3588 (2020).

34. T. B. Jacobs, P. R. LaFayette, R. J. Schmitz, W. A. Parrott, Targeted genome modifications in soybean with CRISPR/Cas9. *BMC Biotechnol.* 15, 16 (2015).

35. T. B. Jacobs, N. Zhang, D. Patel, G. B. Martin, Generation of a Collection of Mutant Tomato Lines Using Pooled CRISPR Libraries. *Plant Physiol.* 174, 2023–2037 (2017).

36. N. Zhang, C. Hecht, X. Sun, Z. Fei, G. B. Martin, Loss of function of the bHLH transcription factor Nrd1 in tomato enhances resistance to Pseudomonas syringae. *Plant Physiol.* 190, 1334–1348 (2022).

37. C. L. M. Gilchrist, Y.-H. Chooi, clinker & clustermap.js: automatic generation of gene cluster comparison figures. *Bioinformatics* 37, 2473–2475 (2021).

38. K.-H. Gartemann, *et al.,* The Genome Sequence of the Tomato-Pathogenic Actinomycete Clavibacter michiganensis subsp. michiganensis NCPPB382 Reveals a Large Island Involved in Pathogenicity ▿ †. *J Bacteriol* 190, 2138–2149 (2008).

39. M. Alexou, A. D. Peuke, Plant Mineral Nutrients, Methods and Protocols. *Methods Mol. Biol.* 953, 195–207 (2012).

40. N. Jiang, *et al.,* Evaluation of suitable reference genes for normalization of quantitative reverse transcription PCR analyses in Clavibacter michiganensis. *Microbiology open* 8, e928 (2019).

41. A. J. Weisberg, *et al.*, A Novel Species-Level Group of Streptomyces Exhibits Variation in Phytopathogenicity Despite Conservation of Virulence Loci. *Mol. Plant-Microbe Interact.* **34**, 39–48 (2021).

42. A. G. Matthysse, S. Stretton, C. Dandie, N. C. McClure, A. E. Goodman, Construction of GFP vectors for use in Gram-negative bacteria other than Escherichia coli. *FEMS Microbiol. Lett.* 145, 87–94 (1996).

43. D. H. Figurski, D. R. Helinski, Replication of an origin-containing derivative of plasmid RK2 dependent on a plasmid function provided in trans. *Proc. Natl. Acad. Sci.*76, 1648–1652 (1979).

44. B. Kessler, V. de Lorenzo, K. N. Timmis, A general system to integrate lacZ fusions into the chromosomes of gram-negative eubacteria: regulation of the Pm promoter of the TOL plasmid studied with all controlling elements in monocopy. *Mol. Gen. Genet. MGG* 233, 293–301 (1992).
